# Shared protocol to enrich and compare biochemical and biophysical properties of Extracellular Vesicles from human plasma and skeletal muscle biopsy

**DOI:** 10.1101/2024.02.19.580950

**Authors:** Valentina Mangolini, Annalisa Radeghieri, Simone Piva, Stefano Cattaneo, Marco Brucale, Francesco Valle, Arianna Balestri, Costanza Montis, Nicola Latronico, Paolo Bergese, Lucia Paolini

**Author notes:** Correspondence to: Dr. Lucia Paolini, Department of Medical and Surgical Specialties, Radiological Sciences and Public Health, University of Brescia. Viale Europa, 11, 25123, Brescia, Italy. e-mail address. ORCID.

## Abstract

**Aim:** Studies on extracellular vesicles (EVs) focused on samples enriched from liquid matrices, such as cell culture media and blood. Recent research highlights the roles of EVs derived from the extracellular matrix of solid tissues, and how investigating these specific EVs offers insights into their microenvironment and potential biological influences on surrounding cells. This study presents a shared method to separate and compare EV enriched from solid (human skeletal muscle biopsy) and liquid (human plasma) matrices, addressing technical challenges and minimizing biases in separation techniques.

**Methods:** Plasma and skeletal muscle-EVs were obtained combining serial centrifugation steps and discontinuous sucrose density gradient. EVs characterization employed advanced analytical techniques such as Western Blot, Colorimetric Nanoplasmonic assay, Atomic Force Miscoscopy, Nanoparticle Tracking Analysis and Dynamic Light Scattering to focus on biomolecular composition, nanomechanical properties, particle yield, size distribution, and colloidal stability.

**Results:** The analysis revealed distinct differences between skeletal muscle-EVs and plasma-EVs including: molecular composition, physical and nanomechanical properties, as well as particle size distribution. Skeletal muscle-EVs exhibited unique colloidal behavior compared to their plasma counterparts, suggesting tissue-specific features that may influence their biological activity and stability.

**Conclusions:** The findings demonstrate that EVs from skeletal muscle tissue possess unique biochemical and biophysical characteristics when compared to those derived from plasma. These differences reflect their diverse biological origins and microenvironments. Understanding these distinctions could advance the development of EV-based diagnostic tools, particularly for muscular disorders, and broaden our knowledge of EV roles across various tissue contexts.

**HIGHLIGHTS:**

- Shared protocol to enrich and compare extracellular vesicles (EVs) from both solid (skeletal muscle biopsy) and liquid (plasma) human samples, reducing methodological bias across matrices.
- Skeletal muscle–derived EVs differ significantly from plasma EVs in biochemical and biophysical properties such as molecular composition, size distribution, nanomechanical properties, and colloidal stability.
- EV heterogeneity across microenvironments emerged as a key biological feature, highlighting their potential for specific diagnostic applications.

## INTRODUCTION

The majority of studies on EVs conducted thus far have mainly focused on EVs separate from liquid matrices such as cell culture media, blood, and urine. This is ascribable to the fact that these samples are readily available and can be processed using established separation techniques^[1,2]^. As a result, our understanding of EVs has been limited to those found in these particular matrices. In the latest years, the important roles of EVs in solid tissues^[3,4]^, including human and non-human brains^[5–10]^, cardiac^[11]^ and skeletal muscles^[12,13,14]^, liver^[14,15]^, adipose tissue^[16,17]^ and several tumors^[18–21]^, are emerging.

The study of solid tissue EVs is expected to offer a glimpse into the EVs in the microenvironment where they are released. Moreover, it provides the potential to comprehend the mechanisms through which EVs, released in the extracellular matrix of a tissue, exert a biological influence on their cellular targets, which may be located in various organs. By investigating solid tissue EVs, we can gain insights into the complex interplay between EVs and their surrounding environment, shedding light on the underlying processes and effects they have on cells.

This increases the possibility to find specific signatures of pathological conditions within organs. By analyzing the distinct profiles of tissue-EV associated with various diseases, researchers can uncover biomarkers that serve as diagnostic indicators. This knowledge could aid in early disease detection but also provides a basis for developing novel liquid biopsy assays^[20,22]^ using blood^[23]^ or other biological fluids^[24]^.

However, this perspective is at its infancy due to technical challenges^[4]^ and the lack of rigorous approaches that allow comparing EVs from solid and liquid tissues, minimizing bias due to different separation techniques and highlighting differences due to the EV microenvironments.

Common techniques applied to separate EVs from liquid matrices imply several combinations of differential (ultra)centrifugation, density gradient, size exclusion chromatography, fluid-flow based separation, affinity-based separation^[1]^ and microfluidics techniques^[25]^. Recently developed EV separation methods from solid tissues mainly involve tissue homogenization or tissue digestion with different enzymes followed by differential (ultra)centrifugation, filtration^[3]^ and density gradients^[4]^. However, it is more and more evident among EV community that “the product is the process”^[26,27]^, indeed the application of different separation methods inevitably leads to the recovery of different EV subpopulations, in terms of physical properties^[28,29]^ (i.e. number density, size distribution, purity from exogenous contaminants), biomolecular content^[30–32]^ (i.e. proteins, metabolites, nucleic acids, biomolecular corona) and therefore biological activity^[29,33]^. In turn separation procedures affect EV functionality both *in vitro*^[34]^ and *in vivo*^[35]^ and this might be due also to the presence of EV co-isolates^[26]^.

Therefore, there is the need of shared EV separation protocols that can be equally applied to liquid and solid matrices in order to compare EVs from these two different sources without the biases that inevitably different separation methods introduce.

This would allow the exploration of how the biomolecular and biophysical properties of EVs may vary between those present in the tissue environment and those circulating, which remains poorly investigated.

To this aim, we have chosen to analyze EVs from human skeletal muscle (SkM) and human plasma derived from blood. These samples have a great clinical relevance: SkM is not only the largest organ in the human body, playing a central role in whole-body energy metabolism, but also acts as a secretory organ, producing and releasing hundreds of products, including myokines and EVs (SkM-EVs) that can have autocrine, paracrine, and endocrine effects^[12,13,36,37]^. Plasma is the most studied source of EVs from bio-fluids^[1,2]^.

Previous studies that compared EVs from plasma and tissues (particularly skeletal muscle) relied on EVs collected from tissues cultured *ex vivo*^[13,38]^. However, this approach introduces new variables related to the culture conditions, meaning that EVs separated *ex vivo* may not accurately represent those directly obtained from the tissue’s extracellular matrix. Another attempt used immunoaffinity-based separation methods^[12]^, hence narrowing the analysis on subpopulations presenting specific surface markers. In addition, these studies have been conducted on animal models, especially rodents, but the translation of those data to human EVs have never been demonstrated.

To minimize modifications induced by differences in the separation process we standardized the procedure to the greatest practicable extent, optimizing the protocol to be applied both to liquid and solid starting materials. This combined method allowed to obtain reproducible and comparable set of EVs from the two different sources. Indeed, SkM-nanoparticles were released from the extracellular matrix in a liquid phase, using mechanical and enzymatic dissociation. EVs from tissue and plasma were, then, enriched using a consistent protocol involving serial ultracentrifugations and discontinuous sucrose gradient allowing to enrich EVs according to their density and size, general properties conserved in different subtypes of EVs.

We, then, characterized the preparations using orthogonal collective and single-particle analysis technique, dissecting sample characteristics at multi-scale levels: from the molecular scale (immunoassays to define their biochemical composition), to the meso-scale (determining samples colloidal properties in terms of purity from exogenous proteins, nano-mechanical features, concentration, size distribution and stability).

This allowed to define “on a common reference” specific biochemical and biophysical properties of EVs secreted by SkM cells and released in the interstitial space^[5,7,12,19]^ in respect to EVs present in plasma, the intercellular matrix of blood. These characteristics have never been described and compared before for EVs from human SkM tissue and plasma.

Studies in this direction might be useful in understanding open questions in EV biology, for example if EVs circulating in the blood stream exhibit different properties than EVs present in the extracellular matrix of different tissue. It has been demonstrated that EVs deriving from healthy or pathological tissues can be detected in circulation^[38–41]^, which raises important considerations about how they enter the bloodstream and whether they need to acquire specific characteristics or undergo changes in their properties after entering circulation.

All this information could help in elucidating the EV roles in various physiological and pathological processes and to develop novel, non-invasive diagnostic and monitoring assays for many diseases including muscular pathologies, hereditary diseases such as metabolic myopathies, dystrophic myopathies or acquired conditions as Intensive-care acquired weakness in critically ill patients with sepsis^[42,43]^.

## METHODS

All data were acquired and described following the MIRABEL^[44]^, MISEV 2018^[2]^ and MISEV2023^[1]^ international guidelines. We have submitted all relevant data of EV samples to the EV-TRACK^[45]^ database (EV-TRACK ID: EV240003) and MIBlood-EV^[46]^ (see Fig.1SI)

**Figure 1.**
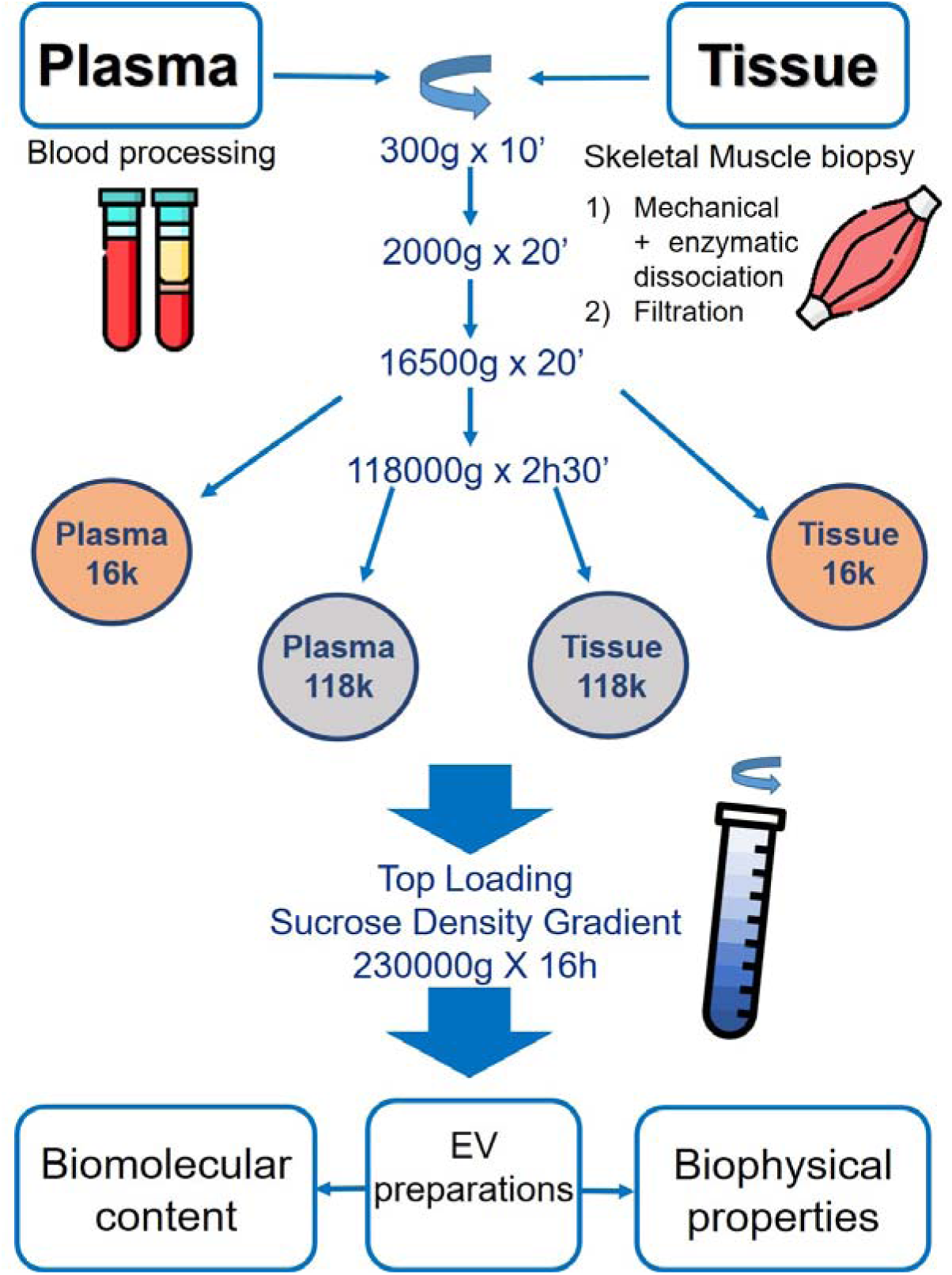
Rationale of the paper sketched. Enrichment and characterization of EVs from human plasma and SkM tissue. After isolation, samples were characterized for their biochemical and biophysically properties. Icons made by Freepik and ITM2101 from www.flaticon.com.

### Subject Recruitment and Sample Collection

In this study, 19 plasma samples and 12 skeletal muscle (SkM) tissue biopsies were obtained from subjects hospitalized in the Intensive Care Unit (n=14) and Orthopedic ward (n=5) of the Spedali Civili University Hospital in Brescia, Italy. The Ethics Committee of Brescia approved the study in adherence with the Declaration of Helsinki and Good Clinical Practice (Protocol number 34 68). All traceable identifiers were removed before analysis to protect patient confidentiality; all samples were analyzed anonymously. Written informed consent was obtained from the patient before the collection of samples.

The study cohort consisted of 19 subjects, including 14 males (73.7%) and 5 females (26.3%), with an age range of 38 – 83 years. (average age 68,2 years). The mean age was 67 years in males and 71,6 years in females. Both blood and SkM tissue biopsies were collected at the same time and processed within 30 minutes from the withdrawal.

Plasma was obtained as previously described from peripheral blood samples in EDTA^[47]^. Blood was collected through a peripheral venous catheter in one of the main venous trunks of the forearm and SkM tissue was collected from quadriceps femoris. Biospecimens were collected and processed, before storage, following international recommendation guidelines^[46,48]^. Samples were stored at -80°C. See Supplementary Materials for more details.

### Enrichment of EVs from human plasma

Plasma EVs were isolated through serial ultra-centrifugation and discontinuous sucrose gradient (DSG). All steps were performed at 4°C. Briefly, 1 ml plasma was centrifuged at 300 x g for 10 min. Supernatant (1 ml) was transferred to a new tube and centrifuged at 2,000 x g for 20 min. Supernatant (1 ml) was transferred to a new tube and centrifuged at 16,500 x g for 20 min and finally supernatant was transferred to an appropriate tube and centrifuged at 118,000 x g for 2h and 30 min. The 16,500 x g centrifugation step allows sedimentation of Plasma 16k sample, while the 118,000 x g ultracentrifugation step allow sedimentation of Plasma 118k sample.

DSG was carried out as follows: Plasma 16k and Plasma 118k pellets, obtained as described above, were further processed adapting the protocols developed in previous studies^[30,47]^. Briefly, Plasma 16k or Plasma 118k samples were re□suspended in 1,000 μl buffer A (10 mM Tris□HCl 250 mM sucrose, pH 7.4), loaded on top of a discontinuous sucrose gradient 15% (600 μl), 20, 25, 30, 40, 60, 65% (400 μl), 70% (800 μl) sucrose in 10 mM Tris□HCl, pH 7.4) and centrifuged at 230,000 x g for 16 h. For Western blot (WB) analysis of fractions 1-12 (250 μl) were precipitated by incubation with 10% trichloracetic acid for 4 h as described in Paolini et al.^[49]^. For all the other analyses, fractions from 6 to 9 (750 μl each) were ultracentrifuged and resupended in 100 μl HPLC water and aliquoted for further analyses. See Supplementary Materials for more details.

### Enrichment of EVs from human SkM tissue

EVs were separated from human SkM tissue after mechanical and enzymatic dissociation, as previously described^[19]^ with some modifications. Separation and subsequent characterizations were performed in accordance with the recommendations from the Solid Tissue Task Force^[48]^ of the International Society for Extracellular Vesicles (ISEV). All steps were performed at 4°C except for the enzymatic digestion. Frozen tissue was thawed 2 min at r.t., washed one with cold PBS 1x and weighted (5 samples, mean 0.89 gr). Tissue was transferred in a 100 mm dish with cold RPMI (medium/tissue ratio: 1ml RPMI/ 0.2 gr tissue) and gently cut into small pieces (2x2x2mm) with a scalpel. Tissue pieces were transferred in a 6-well plate with 0.2 gr of SkM tissue each well. RPMI medium was added to each well to reach 2 ml of total medium. SkM tissue was inoculated with DNase (40U/ml final concentration) and collagenase D (2 mg/ml final concentration) at 37°C for 30 minutes in gently rotation (78 rpm). Medium derived after the enzymatic dissociation was filtered by gravity in a 0,70 µm pore strain in order to eliminate the largest debris. The filtrate was used for EV-enrichment through serial ultra-centrifugation and DSG as described for plasma.

Tissue filtrate was centrifuged at 300 x g for 10 min. Supernatant was transferred to a new tube and centrifuged at 2,000 x g for 20 min. Again, supernatant was transferred to a new tube and centrifuged at 16,500 x g for 20 min in order to obtain Tissue 16K preparation. Finally, supernatant was ultracentrifuged at 118,000 x g for 2 h and 30 min to get Tissue 118K preparation. All centrifugation steps were performed at 4°C.

DSG: Tissue 16k and Tissue 118k pellet obtained were treated as described for plasma preparations. For WB analysis of gradient fractions see section above.

SkM tissue homogenate was obtained from SkM tissue before (THB) and after the mechanical and enzymatic dissociation (TH). See Supplementary Materials for more details.

### SDS-PAGE and WB analysis

SDS−PAGE and WB were performed by standard procedures^[50]^ Sucrose density gradient fractions from 1 to 12 (20 μl of each fraction) were loaded on acrylamide–bisacrylamide, 12.5% gel. Total plasma or tissue homogenate (30 μg, protein content determined by Bradford assay) were loaded in the same gel as positive control of antibody reactivity in Western blot. Samples were electrophoresed and analyzed by WB with the antibodies described in the text. See Supplemental Material for more details.

### COlorimetric NANoplasmonic (CONAN) assay

The EV-enriched preparations (DSG fractions 6-9) were checked for purity from protein contaminants using the COlorimetric NANoplasmonic (CONAN) assay^[51,52]^. EVs were resuspended in 100 μl of HPLC water. Two microlitres of EV solution at serial dilutions in HPLC water (1:1, 2μl of starting sample – 1:10, 2 μl of sample diluted in HPLC H_2_O – 1:100, 2 μl of sample diluted in HPLC water) were resuspended in 23 μl of water, mixed with 50 μl of gold nanoparticles 6 nM and 25 μl of PBS. The result of the assay was collected on Ensight Multi Mode Reader (Perkin Elmer). Measurements were performed for each sample in triplicate.

### Bicinchoninic acid (BCA) and Bradford assay

Protein concentrations of samples (fractions 6-9) were determined with a Pierce™ BCA Protein Assay Kit (ThermoFisher, Rockford, USA), and Bradford assay (Biorad) following the manufacturer’s instructions.

### Atomic Force Microscopy (AFM) analysis

AFM imaging was used to determine sample morphology and quantitative morphometry as described elsewhere^[53,54]^. Five μL aliquots from samples were deposited on glass coverslips previously functionalized with 0.01 mg/mL poly-L-lysine and left to adsorb for 60’ at 4°C. Aliquots were progressively diluted up to 200x in successive depositions in order to maximise the surface density of isolated particles. Imaging was performed in ultrapure water at room temperature on a Bruker Multimode8 AFM equipped with a Nanoscope V controller, a sealed fluid cell and a type JV piezoelectric scanner using Bruker ScanAsystFluid+ probes. Background subtraction was performed using Gwyddion^[55]^ 2.61. Quantitative morphometry was performed with custom Python scripts to recover the surface contact angle (CA) and equivalent solution diameter (D) of several hundred individual objects for each sample. The relative particle concentrations of samples were estimated as described in Supplementary section.

### Nanoparticle Tracking Analysis (NTA)

Pellets from 6 to 9 sucrose fractions obtained from both plasma and human SkM tissue were pooled together, resuspended in 100 µl of sterile HPLC water and for the characterization by Nanoparticles Tracking Analysis (NTA) using a NanoSight NS300 system (Malvern, Panalytical LTD, Malvern, UK) to evaluate the concentration and size distribution of EVs. See Supplemental Material for more details.

### Dynamic light scattering (DLS) analysis and zeta potential measurement

DLS at θ = 90° and zeta potential measurements were performed using a Brookhaven Instrument 90 Plus (Brookhaven Instruments Corporation, Holtsville, NY). For DLS, each measurement was an average of four replicates of 1 minute each. The autocorrelation functions were analysed by Laplace inversion using the CONTIN algorithm, allowing an estimate of the particles size distribution. Zeta potentials were obtained from the electrophoretic mobility u, according to the Helmholtz-Smoluchowski equation: ζ = (η⁄ε) × u where η is the viscosity of the medium and ε is the dielectric permittivity of the dispersing medium. The zeta potential values are given as the average of ten measurements, performed in water.

### Statistical analysis

Significant differences among datasets were determined with Student’s t-test (Graph Pad, OriginLab). P values of less than 0.05 were considered statistically significant with **p< 0.01 ***p < 0.001 and ****p < 0.0001. Values were shown as mean ± standard error of the mean (SEM) of at least 3 experiments.

## RESULTS AND DISCUSSION

### Shared protocol for the enrichment of nanoparticles from human plasma and SkM tissue biopsies

Peripheral blood were collected from 19 subjects and SkM tissue biopsies from 12 subjects. Blood was processed as described in Methods section to obtain plasma. SkM tissue was washed twice in cold solution NaCl 0.9% and sample tube was snap-frozen in liquid nitrogen. Both plasma and SkM tissue were stored at -80°C. Then, Plasma and SkM tissue biopsies were processed with a newly optimized combined protocol (Fig.1).

SkM biopsy (mean weight 0.89 gr) was thawed and washed with cold PBS. The biopsy was then incubated in RPMI medium and processed through gentle mechanical and enzymatic dissociation with DNase and Collagenase for 30 min at 37°C. This process allowed to release nanoparticles embedded in the interstitial space surrounding tissue cells and to diffuse them in the media during the procedure^[19]^. The media containing SkM-EVs was filtered through a 70 μm pore cell strainer in order to eliminate the largest tissue debris. At this stage the same protocol for the EV enrichment from plasma and SkM was used. The SkM filtrate and 1ml of plasma were centrifuged at 300g for 10 min, the supernatant (SN), collected without disturbing the pellet, was centrifuged at 2,000g for 20 min. Then, the subsequent SN was centrifuged at 16,500g for 20 min. The subsequent pellet is referred in the text as the Plasma or Tissue 16k subpopulations. Finally, the SN obtained after the 16,500g centrifugation was centrifuged at 118,000g for 2h 30 min to obtain pellets referred in the text as Plasma or Tissue 118k subpopulations.

It is recognized that pellets derived from both 16k and 118k Plasma and Tissue are composed by a complex mixture of different biogenic nanoparticles^[56]^: extracellular particles (EP- i.e. extracellular vesicles), non-vesicular extracellular particles (NVEP- i.e. lipoproteins, exomeres^[57]^), cellular and tissue small debris. In order to refine the separation of EPs, all four samples were loaded, separately, on top of a discontinuous sucrose gradient (DSG) and centrifuged at 230,000g for 16h. This allows the separation of different nanoparticles both for their size and density^[30,47,58,59]^. Twelve fractions of equal volume (400 μl) were collected, processed as described in Methods section and analyzed as further described.

### Selection of Plasma and Tissue-EV positive fractions by biochemical characterization

Western Blot analysis was conducted for the biochemical characterization of the twelve fractions deriving from the processing of the Plasma 16k, Plasma 118k, Tissue 16k and Tissue 118k pellets by DSG. This allowed to determine the gradient fractions enriched in EVs and fractions enriched in NEVPs markers. Different proteins were selected according to MISEV 2018 guidelines^[2]^ and recent studies^[60]^: EV markers (tetraspannins CD63, CD81; protein involved in the vesicular trafficking such as Flotillin1; single-pass transmembrane protein such as ADAM10; component of the ESCRT□I complex such as TSG101); lipoproteins markers (ApoB marker of Low-density and Very-Low density Lipoproteins -LDL and VLDL; APOA1 marker of High Density Lipoprotein-HDL); SkM cell and SkM-EVs^[12,13]^ markers (Caveolin-3 and Beta-enolase); markers of co-isolates/contaminants^[1,26]^ (hemoglobin A for blood; Fatty Acid binding protein (FABP) for adipose tissue^[61]^; GM130 and Calnexin for intracellular compartments: the Golgi network and the endoplasmic reticulum respectively).

As shown in Fig. 2, fractions 4 and 5 (red box, corresponding to a density of 1.09-1.11 g/cm^3^) resulted enriched in HDL marker ApoA1 both in Plasma 16k (Fig. 2A) and in Plasma 118k (Figure 2B) samples, while ApoB resulted enriched in fraction 4 and 5 for Plasma 118k preparation, and slightly present in fraction 6 and 7 in Plasma 16k preparation. Differences in lipoprotein distribution in gradient fractions could be due to donor-to-donor variations. On the other hand, both Plasma 16k and Plasma 118k fractions from 6 to 9 (corresponding to a density of 1.13-1.29 g/cm^3^) resulted enriched in EV markers, but depleted of blood contaminants such as lipoproteins and hemoglobin A (Supplementary Figure 2-6). Fraction from Plasma samples showed weak signals from markers of the intracellular compartments GM130 and Calnexin, and only in fraction 6 (Fig. 2A, B).

**Figure 2.**
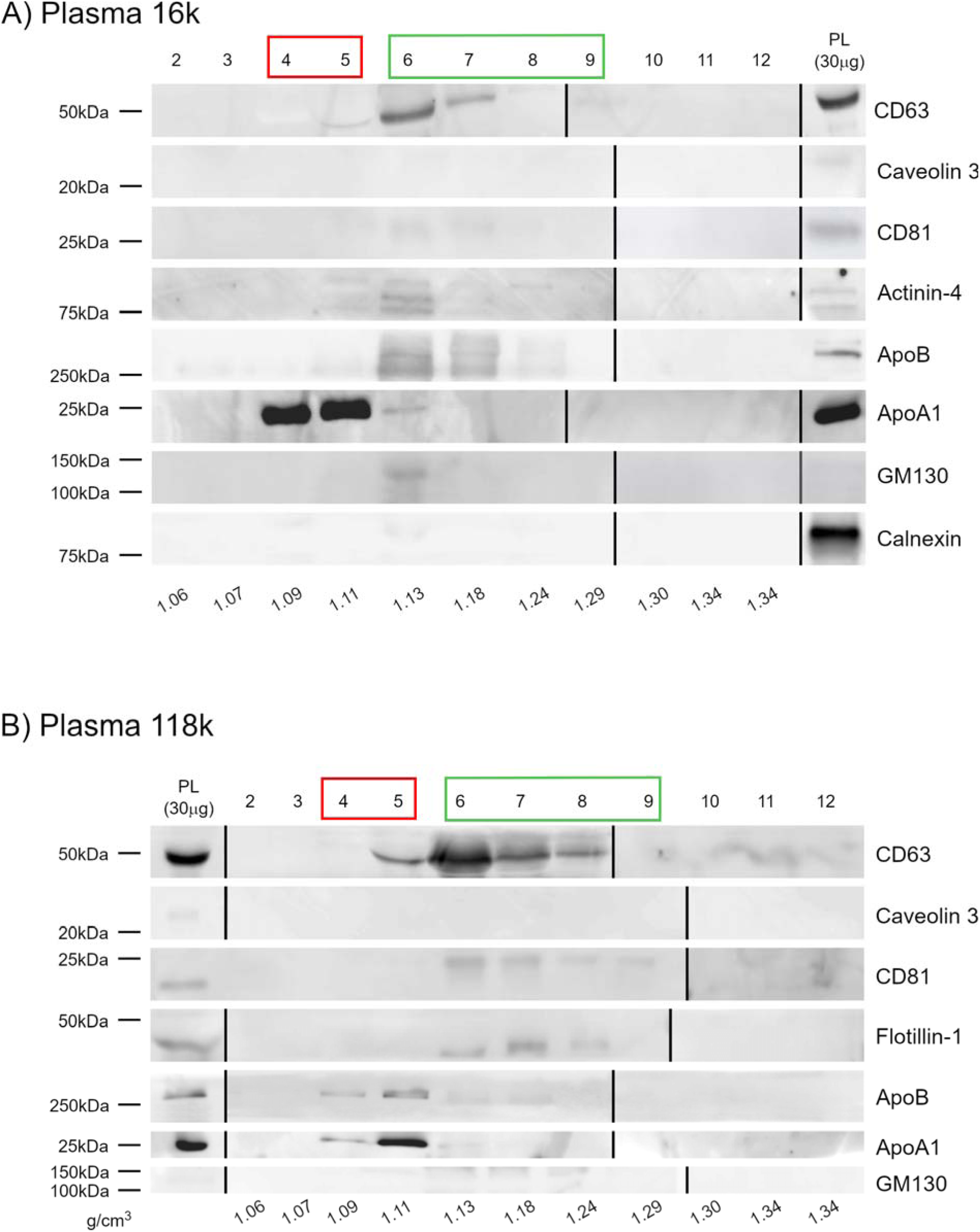
Biochemical characterization of Plasma samples after sucrose gradient fractionation. Equal volume (20 μL) of each fraction both for Plasma 16k (A) and 118k (B) were loaded, 30 μg of Plasma total protein (PL) were loaded as positive control. Samples were electrophoresed on SDS-PAGE gel (12% (Acrylamide/Bis-acrylamide) and analyzed for the antibodies described in the figures. Un-cropped WB are available in the Supplementary Information (Fig.2-5SI). Each fraction density is indicated in g/cm^3^. Red box: fractions discarded for further analyses; Green box: fractions collected and further processed as a single sample.

Similarly, fractions 6 to 9 of Tissue 16k (Fig. 3A) and Tissue 118k (Fig. 3B) samples, after DSG processing, resulted enriched in EV markers such as ADAM10, Flotillin-1, TSG101 and CD81. Fractions analyzed still retain the signal of surface proteins such as TSG101^[62]^, transmembrane proteins such as CD81 and ADAM10^[63]^. These data indicate that the mechanical and enzymatic treatment of the SkM tissue did not alter the EV surface protein profile and we observed an enrichment of these proteins in the EV preparations as expected^[64]^.

**Figure 3.**
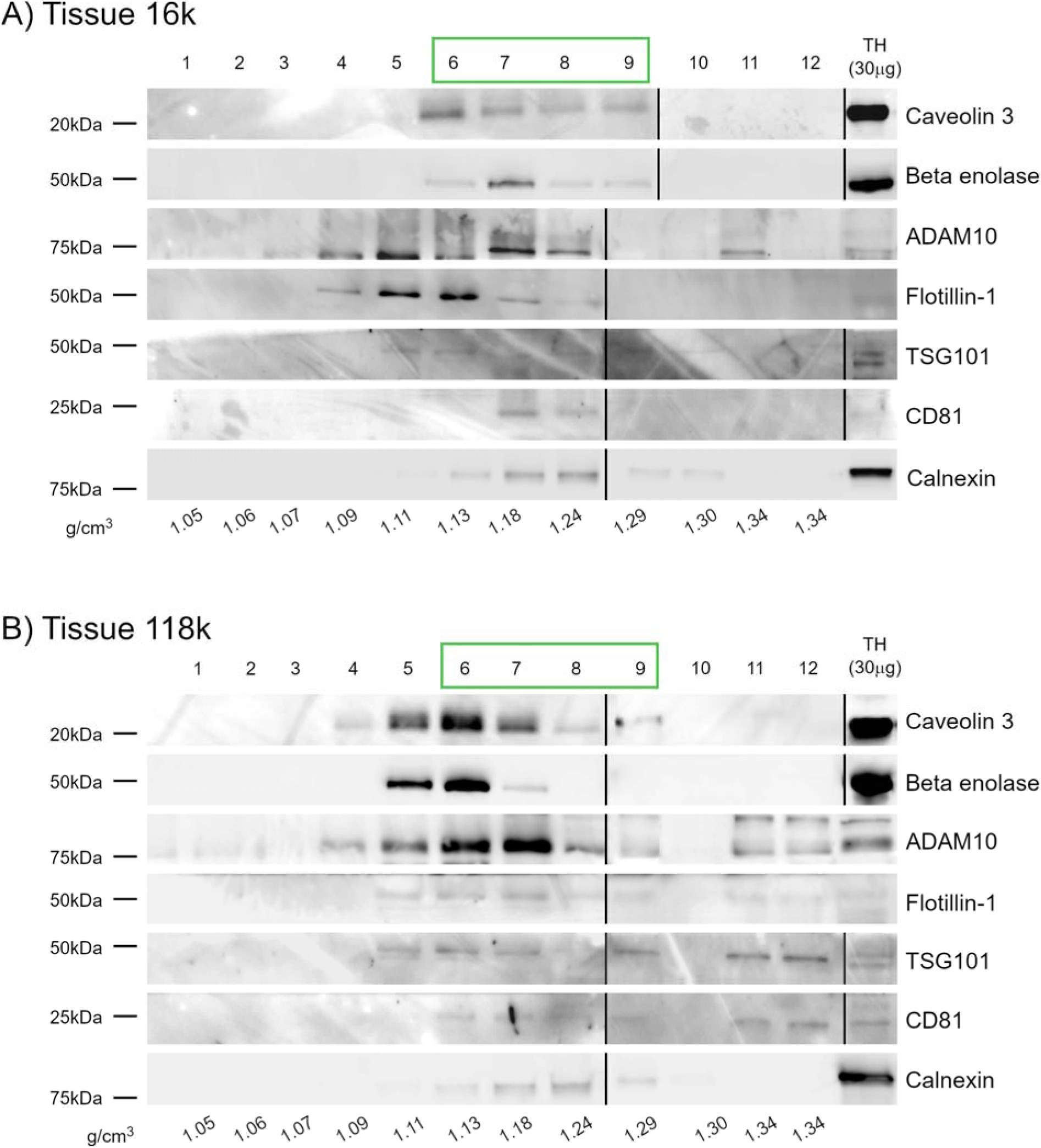
Biochemical characterization of Tissue samples after sucrose gradient fractionation. Equal volume (20 μL) of each fraction both for Tissue 16k (A) and 118k (B), 30 μg of Tissue Homogenate (TH) were loaded as positive controls. WB for the antibodies described in the figures. Un-cropped WB are available in the Supplementary Information (Fig 6-8SI). Each fraction density is indicated in g/cm^3^. The green box indicates fractions that, for further analyses, have been collected and further processed as a single sample.

In addition, fractions 6-9 resulted enriched in SkM cell marker proteins such as Caveolin-3 and beta-enolase indicating the tissue-specific origin of EVs in the preparations, as previously suggested by Watanabe et al^[12]^. Signal intensity of Caveolin-3 in Plasma (Fig. 2A and B, PL) is lower than the signal intensity in Tissue homogenate (Fig. 3A and B, TH) and a similar trend is reflected in gradient fractions from Plasma (Fig.2A,B) and Tissue samples respectively, indicating that, in this experimental conditions, Caveolin-3 could be considered a potential specific SkM-EV marker^[12,14,37]^.

Furthermore, in fraction 6-9 signal of lipoprotein markers and adipose tissue markers were not present (Supplementary Information- SI-, Fig. 6SI,7SI), indicating a negligible blood and adipose tissue contaminants in the SkM tissue biopsies. Tissue samples retain some amount of cellular marker Calnexin^[7,22]^ in fraction 6 to 9, even if in opposite tendency to the signal of EV markers. Quantification of the chemiluminescent signal of Calnexin in SkM tissue homogenate before (THB) and after (TH) enzymatic and mechanical treatment did not change significantly (Fig. 8SI), indicating that the processing did not alter tissue protein composition or induced significant tissue disruption. It is to note that, in our experimental conditions, the chemiluminescent signal of GM130 was not detectable neither in homogenates obtained from the SkM tissue before (data not shown) and after the enzymatic treatment, nor in the gradient fractions, indicating that probably this marker is unsuitable for determining the presence of intracellular co-isolates in human muscle samples, unlike SkM tissue from other species^[12]^ (Fig. 6SI, 7SI).

According to the results described above, we decided to focus our attention to fractions from 6 to 9 (Fig. 2 and 3, green box) for all four preparations: these fractions are enriched in EV markers and contain nanoparticle with the typical density of EVs^[30,47,58,59]^. We discarded fractions 4 and 5 (Fig. 2, 3 red box) that still contain EV markers, but have lower density and, especially in plasma samples, are also enriched in lipoprotein markers. For further analyses fractions 6-9 have been collected and analyzed as a single sample and will be further referred as Plasma 16k fractions 6-9 post gradient (PL16kG), Plasma 118k fractions 6-9 post gradient (PL118kG), Tissue 16k fractions 6-9 post gradient (TS16kG), Tissue 118k fractions 6-9 post gradient (TS118kG).

### Pure Tissue samples from exogenous protein contaminants have a higher protein yield than Plasma Samples

Preparations PL16kG, PL118kG, TS16kG, TS118kG were checked for their purity from exogenous protein contaminants (EPC) with the COlorimetric NANoplasmonic (CONAN) assay as previously described^[51,52,65]^. Serial dilutions (1:1, 1:10 and 1:100 in HPLC water) of the samples were analyzed: PL16kG and PL118kG (Fig. 4A) resulted with an AI% lower than 20% at the minimal dilution of the sample (1:1) and remained below the 20% (dotted line) even at higher dilutions, but with increasing AI%. This represents the trend of samples pure from any detectable EPC (EPC content ≤0.05 μg/μl)^[47,52,66,67]^ where the starting concentration of EVs is optimal for the interaction with gold nanoparticles (AuNPs) and further dilution results in an increase of the AI%.

**Figure 4.**
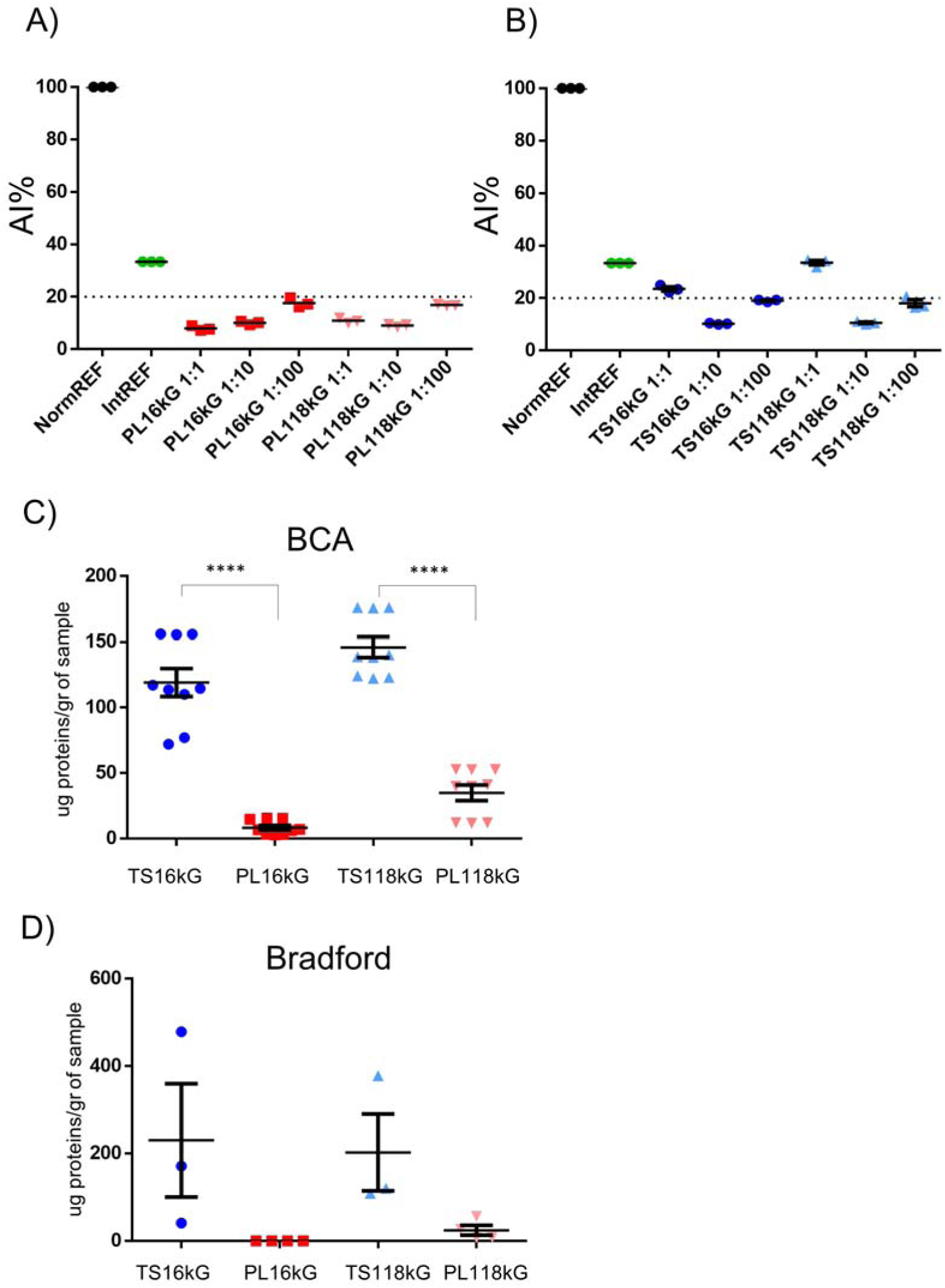
Purity assessment and protein concentration measurements. Fractions 6 to 9 after the sucrose density gradient were collected and analyzed. Samples derived from Plasma (PL16kG, PL118kG) or Tissue (TS16kG, TS118kG). Samples were checked for purity from exogenous protein contaminants (EPC) with the CONAN assay. Aggregation Index ratios (AI%) of serial dilution (1:10, 1:100) of a representative sample are showed in the graph. The dotted line represents the CONAN assay threshold for EPC detection (AI% <20% EPC content ≤0.05 μg/μl). The AI% is inversely proportional to the preparation purity and samples with AI% below the 20% are considered pure. AI% values of a representative sample with assay internal controls (NormREF, black dots; IntREF, green dots) (A); Protein concentration with BCA (B) and Bradford assay (C); Protein concentration for each group was normalized by tissue mass (per 1 gr). Values were shown as mean ± SEM. Significant differences among datasets were determined with Student’s t-test. P values: ****p < 0.0001. TS16kG, dark blue points; PL16kG, dark red squares; TS118kG, light triangle up; PL118kG, light red triangle down.

TS16kG and TS118kG (Fig.4B) presented an AI% higher than 20% at the dilution 1:1, while it decreased under the 20% with the serial dilution 1:10 and 1:100, meaning that the samples are pure from EPC, but a further dilution is needed to fall the AI below the 20% dotted line. These data indicate that our experimental procedures allowed to obtain EVs from SkM tissue pure from EPC, and indicate that both Tissue-EV preparations are more concentrated than Plasma-EV preparations, since TS16kG and TS118kG reached an AI% below 20% after a dilution 1:10 of the starting material. The increase of AI% in further dilutions indicates that samples reached the optimal EV concentration for AuNPs interaction at dilution 1:10, while at dilution 1:100 we are shifting from the optimal range in terms of EV concentration to interact with AuNPs resulting in an increase of the AI%.

After the purity check with the CONAN assay we performed Bicinchoninic acid (BCA) and Bradford assays to determine the protein concentrations in the samples (very likely associated to EVs). In order to compare the protein yield of samples from SkM tissue and plasma, the concentration was normalized to one gram of starting material^[7,19,38,68]^ (1 ml of Plasma and 1 gr of Tissue). BCA and Bradford results were comparable and in accordance to CONAN results. Tissue samples presented a significant higher protein yield than Plasma samples both with 16kG and 118kG preparations measured with BCA (Fig. 4C). It is to note that for the Bradford assay, these highly purified with low-yield samples challenged the assay sensitivity. Consequently, the protein concentration exhibited wide Standard Error of the Mean (SEM), especially in TS16kG and TS118kG, and in the case of the PL16kG sample, it was undetectable (Fig. 4D). However, despite these challenges, the trend of higher protein yield in Tissue samples compared to Plasma samples persisted, as indicated by BCA measurements.

### Tissue and Plasma EV-enriched fractions display heterogeneous nanomechanical properties

The morphology and nanomechanical characteristics of nanoparticles present in PL16kG, PL118kG, TS16kG and TS118kG preparations were evaluated using a high-throughput AFM imaging approach described previously^[53,69]^. Preliminary qualitative inspection of the resulting Atomic Force Microscopy (AFM) micrographs revealed that all samples contained abundant globular particles with a morphology broadly compatible with that typically displayed by EVs (Fig. 5A,B,C,D). This indicates that in all samples analyzed, EVs maintain their morphological integrity after the enrichment protocols.

**Figure 5.**
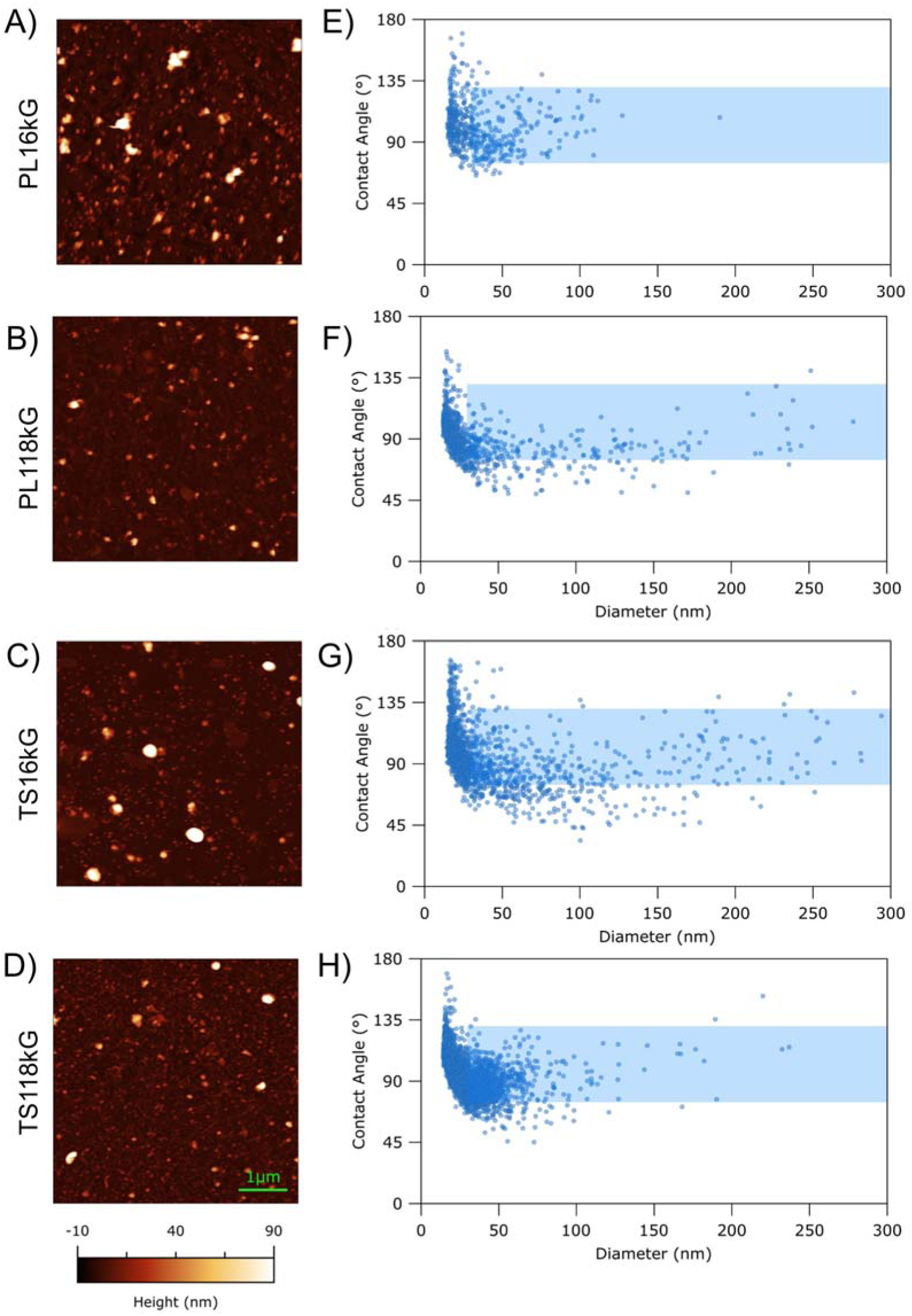
Nano-mechanical features of samples from Plasma and Tissue. Representative AFM images of PL16kG (A), PL118kG (B), TS16kG (C), and TS118kG (D). Contact angle VS diameter plot of several hundred individual particles from at least five different depositions for each sample as calculated via AFM morphometry of PL16kG (E), PL118kG (F), TS16kG (G) and TS118kG (H). Blue box represents area populated by structures with the typical EV-like characteristics: contact angle ranging from 75° to 135°, diameter from 30 to 300 nm.

We recently demonstrated that quantitative AFM morphometry can discriminate different subpopulations of nanoparticles in the same preparation on the basis of their nanomechanical properties. In particular, the stiffness of individual putative vesicles adhered to the substrate can be estimated via their equivalent spherical cap contact angle (CA). In samples derived from plasma, this technique was shown to discriminate EVs from NVEPs such as lipoproteins^[54]^. Briefly, since CAs were shown to be heavily influenced by the nanomechanical properties of their parent particle (*e.g.* the stiffness of intact EVs), collecting the morphometric measurements of individual objects from the same sample on a CA versus Diameter (D) plot results in the clustering of different subtypes of EVs and NVPs in specific zones of the plot^[54]^.

Accordingly, the CA and D values of several hundred individual particles from PL16kG, PL118kG, TS16kG and TS118kG preparations were collected to evaluate the particle heterogeneity of each sample (Fig.5). All samples exhibited the typical log-normal distribution of sizes, containing abundant globular particles with D values below the range typically associated with intact EVs (<30 nm) and highly dispersed CA values surpassing those previously associated with lipoproteins (> 135°); although it is impossible to determine the biological identity of these particles via AFM, their CA and D ranges might be compatible with protein aggregates and/or other NVEPs. Given their higher stiffness compared to liposomes, they could be protein-enriched NEVPs, similar to exomeres^[57]^.

Their overall similarity notwithstanding, the main differences between the four samples are found in the relative abundance, size and stiffness of particles with diameters above 50 nm. As shown in Fig. 5E, PL16kG was found to contain a large fraction of particles within typical CA and D ranges previously associated with intact EVs from a variety of sources^[53,54,69]^ (blue box, CA=80°-130° and D>30nm). On the other hands, the regions with lower CA (<80 nm), ascribed to other biogenic nanoparticles such as lipoproteins, was practically devoid of particles, confirming results from WB. The corresponding PL118kG preparation (Fig. 5F) also contained a rich population of EV-compatible particles and a marginal presence of particles with lower CA values. We then applied the same analytical procedure to TS16kG (Fig. 5G) and TS118kG (Fig. 5H) samples. Despite being very similar to the Plasma preparations with abundant particles within typical CA and D ranges previously associated with intact EVs (blue box), both TS16kG and TS118kG samples showed a higher abundance of softer particles (CA below 80°, below the blue box) in the fraction of sizes above 50 nm. This means that the EV subpopulations in the samples enriched from SkM tissue present a more heterogeneous CA than EVs enriched from Plasma.

In addition, as outlined in the methods section, particle surface density calculations from AFM images allow estimating relative particle concentrations across different samples. With this method, the concentration of EVs in samples enriched from tissue (TS16kG and TS118kG) was found to be on average 10^2^ times higher than the corresponding preparations from PL16kG and PL118kG as suggested by CONAN, BCA and Bradford assays (SI, Table 1).

### Tissue samples have higher particles concentration, broader size distribution and higher colloidal stability than Plasma samples

The analytical techniques described earlier provide an estimation of whether one sample is more concentrated than another in terms of nanoparticles and proteins. To confirm these data, we compared the particle concentration in the samples by Nanoparticle Tracking Analysis (NTA). Indeed, NTA results confirmed the data from CONAN, protein quantification and AFM, indicating a significant higher particle yield in TS16kG samples (mean concentration 9,31E+11 particles/gr of tissue) than PL16kG samples (mean concentration 4,14E+10 particles/gr of plasma) and TS118kG (mean concentration 1,43E+12 particles/gr of tissue) samples respect to PL118kG samples (mean concentration 6,57E+10 particles/gr of plasma). It is to note that, to facilitate the comparison of particle yield between Plasma and Tissue samples, data have been normalized to one gram of starting material (Fig. 6A). Despite the differences in nature of the two, in our experimental conditions, all the orthogonal techniques implied confirmed that the EV yield from SkM is significantly higher than Plasma for both subpopulations analyzed.

**Figure 6.**
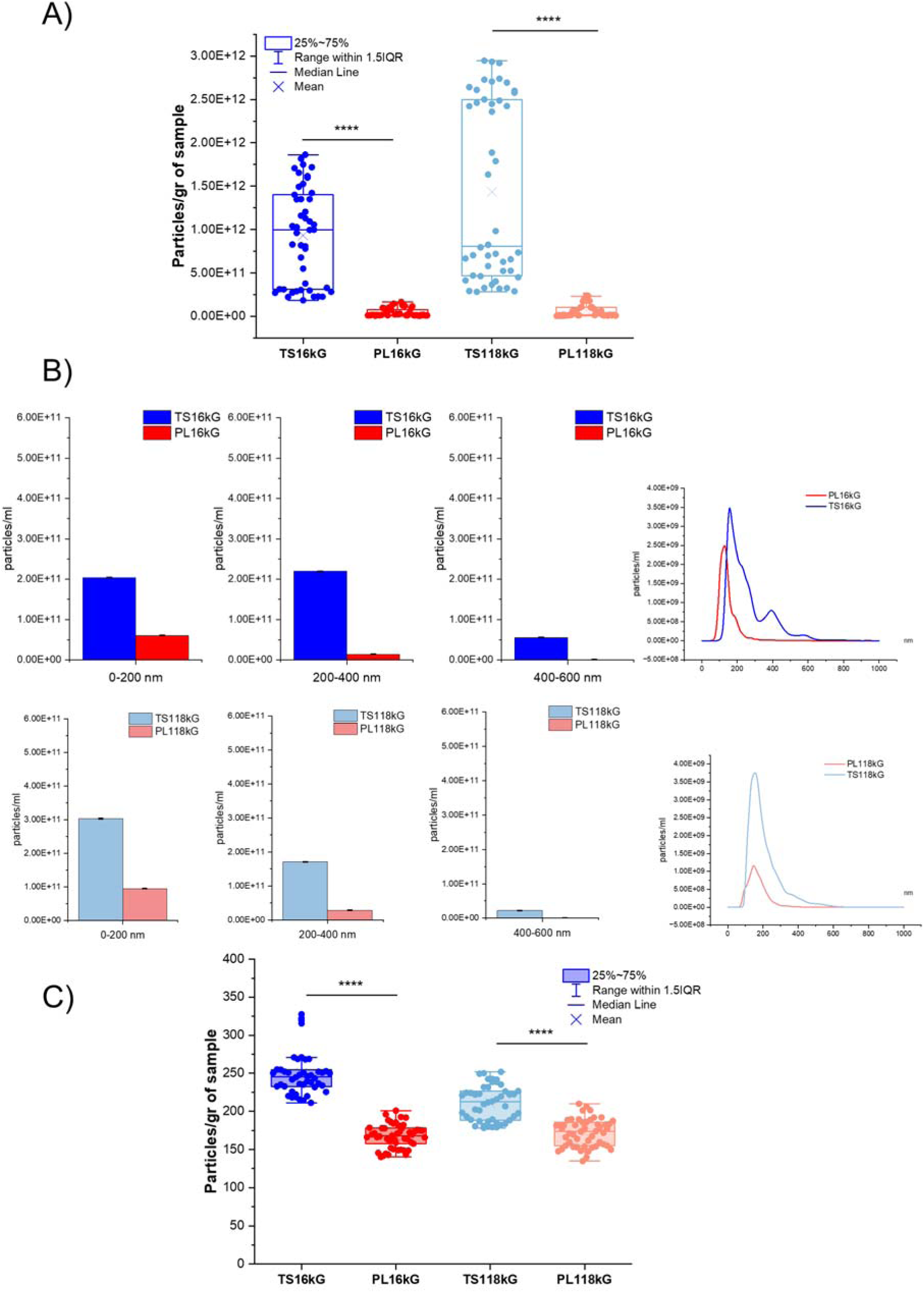
Concentration and size distribution. Particle concentration for each sample were measured by NTA. Particle concentration was normalized for starting material mass (1gr). Box plots represent the interquartile range (25th–75th percentile), with whiskers extending to 1.5×IQR; the central line indicates the median and the “X” denotes the mean. (A). Particle Size distribution was measured by NTA. Histograms represent the mean of the size distribution divided by range (30-200 nm; 201-400 nm; 401-600 nm) of the 16k and 118k samples from plasma and tissue B). Particle size was analyzed by NTA and meanmodal distribution (nm) (C). Data are displayed as box plots (25th–75th percentile, whiskers at 1.5×IQR), with median (line) and mean (X). Significant differences among datasets were determined with Student’s t-test. P values: **p< 0.01, ***p < 0.001 and ****p < 0.0001.

Samples size distribution was determined with orthogonal techniques: NTA, Dynamic Light Scattering (DLS) and AFM. Each method has its advantages and limits and the combined evaluation of data obtained with different approaches allows to reveals different aspect and characteristics.

The size of particles was analyzed using NTA and mean size distribution profiles of preparations are shown in Fig. 6B. According to these measurements, both TS16kG and TS118kG samples have a broader size distribution than PL16kG and PL118kG respectively, with presence of EVs in the range from 30-200 nm, from 201 to 400 nm and also from 401 to 600 nm, while the majority of EVs from Plasma samples are in the range from 30-200 and 201-400 nm (Figure 6B). Notably, TS16kG samples showed a bimodal size distribution, indicating the presence of two main EV populations.

The mean size was found to be significantly different from TS16kG and PL16kG samples, with TS16kG EVs diameter larger than PL16kG EVs (249,70 ± 4,39 nm and 167,42 ± 2,21 nm respectively, Fig. 6C). A significant difference was also observed between TS118kG and PL118kG preparations (210,74 ± 3,41 nm and 171,84 ± 2,56 nm respectively, Fig. 6C). NTA results data indicated that in TS16kG preparation the mode size of EVs is significantly higher than TS118kG sample. We also analyzed NTA data to compare the particle concentration and size distribution of EVs enriched from TS and PL fractions obtained at 16kG and 118kG.

TS16kG and TS118kG preparations significantly differ for concentration and size. In particular, the higher particle counts has been observed in the TS118kG which present the smaller mean size (Fig.9A SI).

On the other hand, PL16kG presented significant lower concentration respect to PL118kG preparations while no significant differences were detected in size (Fig.9B SI).

The Laplace inversion analysis using the CONTIN algorithm was applied to autocorrelation functions obtained through DLS for four preparations (PL16kG, PL118kG, TS16kG, and TS118kG, Fig. 7A,D,G,L) to qualitatively confirm the presence of submicrometer particles and possible aggregates that may be present in EV samples^[70]^. The DLS profiles show a polydisperse size distribution, likely arising from a combination of the scattering from single and aggregated EVs, as well as from the scattering from objects of similar refractive index (e.g., proteins). The scattered intensity-weighted size distribution, depicted in Fig. 7B,E,H,M, provides a reliable analysis without assuming the composition of different populations. The results indicated that PL16kG and TS16kG samples are characterized by the presence of objects ranging from hundreds of nanometers to micrometers in diameter. In contrast, PL118kG and TS118kG preparations exhibit bi-modal size distributions with a small subpopulation (around 100 nm) and larger aggregates (diameter > 400 nm). It should be noted that DLS technique inherently tends to overestimate the presence of larger objects compared to smaller ones^[70]^. Therefore, a number-weighted distribution is also presented (Fig. 7C,F,I,N). In this scale, the majority of the scattering objects are around 100 nm diameter.

**Figure 7.**
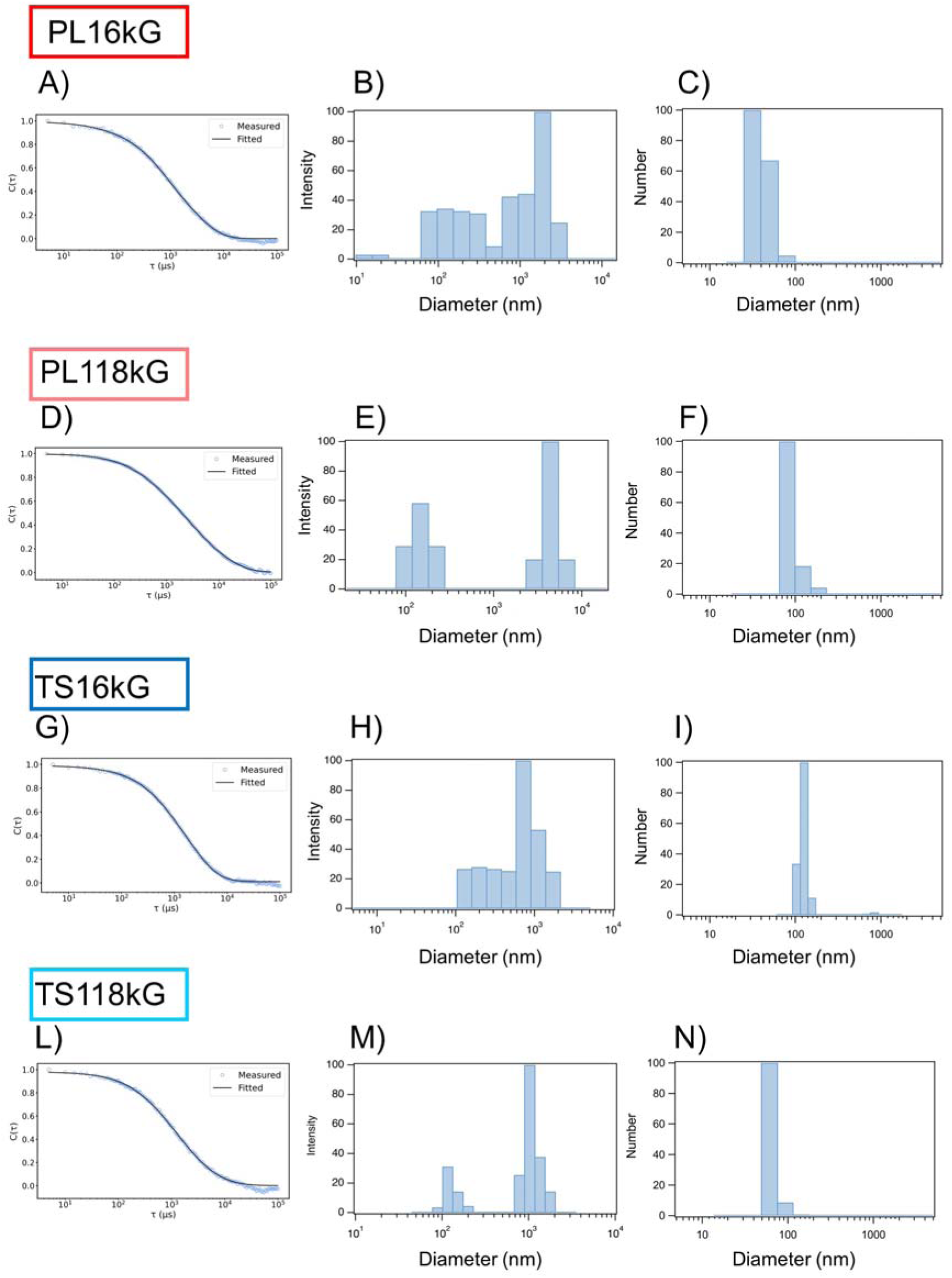
Size distribution with DLS. Representative autocorrelation function of the scattered intensity, with the fitting curve according to the Laplace Inversion via CONTIN algorithm (A,D,G,L); corresponding scattered intensity-weighted (B,E,H,M) and number-weighted (C,F,I,N) size distributions of scattering objects of samples described in the text.

This observation is consistent with the presence of small objects such as EVs, as corroborated by results from NTA and AFM and confirmed the vesicular structure preservation after the separating procedures.

Size distributions obtained via AFM morphometry can be inferred from the abundance of individual particles along the horizontal (D) axis of plots in Fig. 5E-5H. While a direct quantitative comparison of sizes distributions obtained from AFM and NTA is often questionable, the general trends observed by both techniques coincide: both AFM and NTA attribute to TS16kG EVs a higher diameter mode than PL16kG EVs (Fig. 5G and 5E, respectively), while those of TS118kG and PL118kG did not differ significantly (Fig. 5H and 5F, respectively). AFM also attributes to TS16kG EVs (Fig. 5G, blue box) a more heterogeneous size distribution than PL16kG EVs (Fig. 5E, blue box), with more particles populating the area of diameter between 100 and 300 nm in TS16kG sample, while PL118kG and TS118kG EVs (Fig. 5F and 5H, respectively) seem to be more homogenous.

Finally, we analyzed the colloidal stability of the four preparations measuring the zeta-potential. All measurements were done in milliQ H_2_0. TS16kG presented a mean zeta potential of -34.2 mV, PL16kG -13.7mV, TS118kG -24.2 and PL118G -10.6mV (Fig. 8). Notably, differences in colloidal stability between Plasma and Tissue-EVs enriched samples emerged. Both PL16kG and PL118kG exhibited a significantly higher zeta-potential compared to TS16kG and TS118kG, suggesting lower stability. This disparity implies that plasma EVs may be more susceptible to interactions with other components circulating in blood (i.e. other biogenic entities such as lipoproteins^[71]^) than EVs in the extracellular matrix of SkM tissue. It will be interesting to evaluate if this property is specific of blood EVs at the moment of the secretion or if EVs acquired it during the stay in a such a complex biofluid, and if these differences affect the biological functionality of EVs. For example, plasma EVs, with their lower stability, might be more prone to aggregation or faster clearance from circulation, whereas more stable tissue EVs may persist longer in the interstitial space. Tissue EVs, on the other hand, maybe more stable in their local environment due to less interaction with foreign biogenic components. Further analyses should be conducted to assess how would these influence the development of EV-based diagnostics, particularly in distinguishing between tissue and blood-derived EVs, as stability could influences EV bioavailability and detectability in biofluids^[72]^.

**Figure 8.**
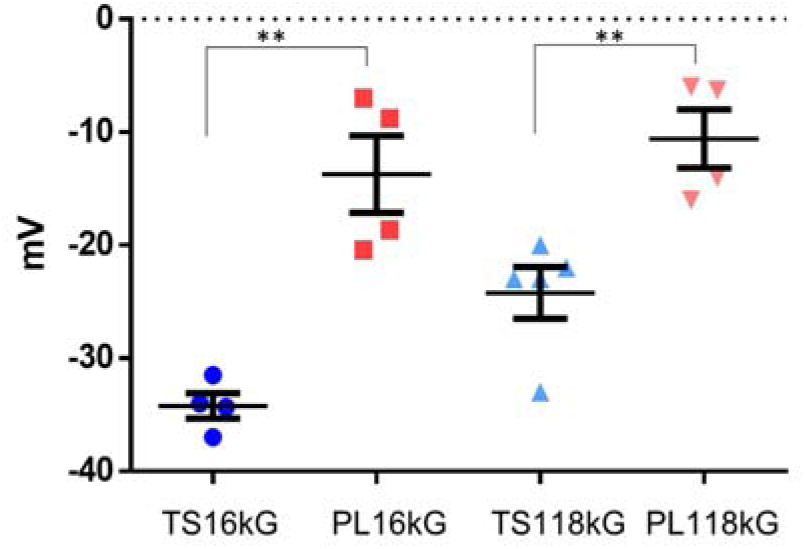
Zeta-potential measurement. Preparations were analyzed for their colloidal stability measuring z-potential with DLS. Values in mV were shown as mean ± standard error of the mean (SEM). Significant differences among datasets were determined with Student’s t-test. P values: **p< 0.01.

## CONCLUSIONS

We presented a shared method to separate EV from liquid and solid matrices, that allowed to compared, for the first time, properties of EVs from human plasma and skeletal muscle tissue. The combination of differential (ultra)centrifugation steps and a top-down sucrose density gradient allowed the separation of EVs from their starting materials and to select gradient fractions that were positive for the presence of typical EV markers and cellular origin markers from SkM tissue. In addition, we demonstrated, with orthogonal techniques, that the selected gradient fractions are depleted from co-isolated NVEPs such as lipoproteins, and contaminants such as cellular debris and exogenous soluble proteins.

Furthermore, the characterization of the preparations showed differences in biomolecular and biophysical properties of EVs-enriched samples from human plasma and SkM. The observed differences regarding the nanomechanical properties, particle yield, size and colloidal stability are not biased by the separation method and thus can reflect specific characteristics of EVs-enriched samples from liquid and solid matrices. This poses new issues in understanding the determinants of these differences: the cellular origin, the molecular composition, the complex interaction between EVs and the biological environment. The last point highlights the importance of considering, in this scenario, the role of the biomolecular corona^[65,73]^ the extravesicular cargo of molecules adsorbed to EVs, influenced by the surrounding matrix and that can change their physicochemical properties^[74]^. In addition, it will be interesting to compare the biophysical properties of EVs from different human tissues enriched with the same protocol to see if they are SkM tissue-specific or typical of all tissue derived EVs.

From a broader standpoint, our optimized method could be widely adopted in the EV field since it can be easily translated to other tissues. In this way, researchers may directly assess the traits that we highlighted to other solid and liquid matrices such as tumor tissues, urines and determine novel pivotal factors in the discovery of biomarkers^[75]^. Infact, the characterization of EV biophysical properties could serve, in the next future, as powerful diagnostic tools^[74,75]^ for various pathological conditions, including, but not only related to SkM pathologies, complementing traditional biomolecular markers.

We are aware that many pre-analytical variables such as storage conditions and freeze-thaw process could influence EVs quality, composition, and function both from solid and liquid matrices.

Practical limitations, including feasibility, logistics and accessibility to fresh tissue samples, often necessitate alternative approaches^[76]^. Currently, solid tissue EVs are most commonly extracted from human and mouse post-mortem tissue that has been fresh frozen at -80°C. However, following international recommendations handling and preservation of human biospecimens we tried to obtain the best quality EVs possible in terms of integrity and purity^[77,78]^.

It has been previously demonstrated that tissue storage at -80°C could slight upregulate cell-associated markers, a result that was consistent with the suggestion that cell rupture may cause intracellular vesicles to overflow. However, other phenotypic characteristics of EVs, such as their purity and morphology, were not significantly altered^[79,80]^, as confirmed from our data.

In conclusion, our results obtained from stored tissue can be of interest for all researchers that can have access to existing biobanks thus increasing the number of samples that can be processed and used as source of EVs, and greatly increasing the possibility for developing EV-based diagnostic tools.

## DECLARATIONS

## Acknowledgements

We thank Dr. Federica Bono from the Dept. of Molecular and Translational Medicine of the University of Brescia for the Ultra-Turrax T25 basic (IKA-Werke) homogenizer.

## Ethical Approval and Consent to Participate

Research was conducted in accordance in accordance with The Code of Ethics of the World Medical Association (Declaration of Helsinki) and Good Clinical Practice. The study was approved by the ethics committee of Spedali Civili di Brescia (Protocol number 3468). Informed consent was obtained from all the subjects enrolled in the study. The privacy rights of subjects involved in the study have always been observed.

## Consent for Publication

All authors have reviewed and endorsed this version of the manuscript for publication.

## Availability of Data and Materials

The CONAN and DLS datasets supporting the conclusions of this article are available in the Zenodo repository, https://doi.org/10.5281/zenodo.10579305. All other data are available on request.

## Authors’ Contributions

CRediT taxonomy. VM Data curation, Formal analysis, Investigation, Methodology, Writing - review & editing; AR Conceptualization, Data curation, Formal analysis, Investigation, Funding acquisition, Writing - review & editing; SP Conceptualization, Data curation, Formal analysis, Funding acquisition, Writing - review & editing; SC Investigation, Methodology, Writing - review & editing; NL Conceptualization; Data curation; Formal analysis; Funding acquisition, Writing - review & editing; MB Investigation, Methodology, Data curation, Writing - review & editing; FV Investigation, Methodology, Data curation; AB Investigation, Methodology, Writing - review & editing; CM Investigation, Methodology, Writing - review & editing; PB Conceptualization; Data curation; Formal analysis; Funding acquisition, Writing - review & editing; LP Conceptualization, Data curation, Formal analysis, Funding acquisition, Investigation; Methodology, Writing - original draft

## Declaration of generative AI and AI-assisted technologies in the manuscript preparation process

During the preparation of this work the author(s) used ChatGPT in order to check English grammar and improving language clarity during manuscript preparation. After using this tool/service, the author(s) reviewed and edited the content as needed and take(s) full responsibility for the content of the published article.

## Supplementary Materials

The datasets supporting the conclusions of this article are included within the article and its additional file named “Supplementary Materials”.

## REFERENCES

1. Welsh JA, Goberdhan DCI, O’Driscoll L, et al. Minimal information for studies of extracellular vesicles (MISEV2023): From basic to advanced approaches. J Extracell Vesicles 2024;13:e12404. [PMID: 38326288 DOI: 10.1002/jev2.12404].

2. Théry C, Witwer KW, Aikawa E, et al. Minimal information for studies of extracellular vesicles 2018 (MISEV2018): a position statement of the International Society for Extracellular Vesicles and update of the MISEV2014 guidelines. J Extracell Vesicles 2018;7:1535750. [PMID: 30637094 DOI: 10.1080/20013078.2018.1535750].

3. Li S-R, Man Q-W, Gao X, et al. Tissue-derived extracellular vesicles in cancers and non-cancer diseases: Present and future. Journal of Extracellular Vesicles 2021;10:e12175. [DOI: 10.1002/jev2.12175].

4. Zhi Z, Sun Q, Tang W. Research advances and challenges in tissue-derived extracellular vesicles. Frontiers in Molecular Biosciences 2022;9.

5. Perez-Gonzalez R, Gauthier SA, Kumar A, Levy E. The Exosome Secretory Pathway Transports Amyloid Precursor Protein Carboxyl-terminal Fragments from the Cell into the Brain Extracellular Space*. Journal of Biological Chemistry 2012;287:43108–15. [DOI: 10.1074/jbc.M112.404467].

6. Vella LJ, Scicluna BJ, Cheng L, et al. A rigorous method to enrich for exosomes from brain tissue. Journal of Extracellular Vesicles 2017;6:1348885. [DOI: 10.1080/20013078.2017.1348885].

7. Huang Y, Cheng L, Turchinovich A, et al. Influence of species and processing parameters on recovery and content of brain tissue-derived extracellular vesicles. Journal of Extracellular Vesicles 2020;9:1785746. [DOI: 10.1080/20013078.2020.1785746].

8. Brenna S, Altmeppen HC, Mohammadi B, et al. Characterization of brain-derived extracellular vesicles reveals changes in cellular origin after stroke and enrichment of the prion protein with a potential role in cellular uptake. J Extracell Vesicles 2020;9:1809065. [PMID: 32944194 DOI: 10.1080/20013078.2020.1809065].

9. Zhang Z, Yu K, You Y, et al. Comprehensive characterization of human brain-derived extracellular vesicles using multiple isolation methods: Implications for diagnostic and therapeutic applications. Journal of Extracellular Vesicles 2023;12:12358. [DOI: 10.1002/jev2.12358].

10. Gomes PA, Bodo C, Nogueras-Ortiz C, et al. A novel isolation method for spontaneously released extracellular vesicles from brain tissue and its implications for stress-driven brain pathology. Cell Communication and Signaling 2023;21:35. [DOI: 10.1186/s12964-023-01045-z].

11. Liang W, Najor RH, Gustafsson ÅB. Protocol to separate small and large extracellular vesicles from mouse and human cardiac tissues. STAR Protocols 2024;5:102914. [DOI: 10.1016/j.xpro.2024.102914].

12. Watanabe S, Sudo Y, Makino T, et al. Skeletal muscle releases extracellular vesicles with distinct protein and microRNA signatures that function in the muscle microenvironment. PNAS Nexus 2022;1:pgac173. [PMID: 36714847 DOI: 10.1093/pnasnexus/pgac173].

13. Ismaeel A, Van Pelt DW, Hettinger ZR, et al. Extracellular vesicle distribution and localization in skeletal muscle at rest and following disuse atrophy. Skeletal Muscle 2023;13:6. [DOI: 10.1186/s13395-023-00315-1].

14. Matejovič A, Wakao S, Kitada M, Kushida Y, Dezawa M. Comparison of separation methods for tissue-derived extracellular vesicles in the liver, heart, and skeletal muscle. FEBS Open Bio 2021;11:482–93. [PMID: 33410274 DOI: 10.1002/2211-5463.13075].

15. Olofsson Bagge R, Berndtsson J, Urzì O, Lötvall J, Micaroni M, Crescitelli R. Three-dimensional reconstruction of interstitial extracellular vesicles in human liver as determined by electron tomography. J Extracell Vesicles 2023;12:e12380. [PMID: 38010190 DOI: 10.1002/jev2.12380].

16. Jeurissen S, Vergauwen G, Van Deun J, et al. The isolation of morphologically intact and biologically active extracellular vesicles from the secretome of cancer-associated adipose tissue. Cell Adh Migr 2017;11:196–204. [PMID: 28146372 DOI: 10.1080/19336918.2017.1279784].

17. Crewe C, Joffin N, Rutkowski JM, et al. An Endothelial-to-Adipocyte Extracellular Vesicle Axis Governed by Metabolic State. Cell 2018;175:695–708.e13. [DOI: 10.1016/j.cell.2018.09.005].

18. Hurwitz SN, Olcese JM, Meckes DG. Extraction of Extracellular Vesicles from Whole Tissue. J Vis Exp 2019. [PMID: 30799860 DOI: 10.3791/59143].

19. Crescitelli R, Lässer C, Lötvall J. Isolation and characterization of extracellular vesicle subpopulations from tissues. Nat Protoc 2021;16:1548–80. [PMID: 33495626 DOI: 10.1038/s41596-020-00466-1].

20. Jang SC, Crescitelli R, Cvjetkovic A, et al. Mitochondrial protein enriched extracellular vesicles discovered in human melanoma tissues can be detected in patient plasma. J Extracell Vesicles 2019;8:1635420. [PMID: 31497264 DOI: 10.1080/20013078.2019.1635420].

21. Tassinari S, D’Angelo E, Caicci F, et al. Profile of matrix-entrapped extracellular vesicles of microenvironmental and infiltrating cell origin in decellularized colorectal cancer and adjacent mucosa. Journal of Extracellular Biology 2024;3:e144. [DOI: 10.1002/jex2.144].

22. Crescitelli R, Filges S, Karimi N, et al. Extracellular vesicle DNA from human melanoma tissues contains cancer-specific mutations. Frontiers in Cell and Developmental Biology 2022;10.

23. Cheng L, Vella LJ, Barnham KJ, McLean C, Masters CL, Hill AF. Small RNA fingerprinting of Alzheimer’s disease frontal cortex extracellular vesicles and their comparison with peripheral extracellular vesicles. Journal of Extracellular Vesicles 2020;9:1766822. [DOI: 10.1080/20013078.2020.1766822].

24. Dong L, Feng M, Kuczler MD, et al. Tumour tissue-derived small extracellular vesicles reflect molecular subtypes of bladder cancer. Journal of Extracellular Vesicles 2024;13:e12402. [DOI: 10.1002/jev2.12402].

25. Abreu CM, Costa-Silva B, Reis RL, Kundu SC, Caballero D. Microfluidic platforms for extracellular vesicle isolation, analysis and therapy in cancer. Lab Chip 2022;22:1093–125. [DOI: 10.1039/D2LC00006G].

26. LargeCscale production of extracellular vesicles: Report on the “massivEVs” ISEV workshop - Paolini - 2022 - Journal of Extracellular Biology - Wiley Online Library. Available from: https://isevjournals.onlinelibrary.wiley.com/doi/full/10.1002/jex2.63. [Last accessed on January 29, 2024].

27. Buschmann D, Mussack V, Byrd JB. Separation, characterization, and standardization of extracellular vesicles for drug delivery applications. Advanced Drug Delivery Reviews 2021;174:348–68. [DOI: 10.1016/j.addr.2021.04.027].

28. Jeon H, Kang S-K, Lee M-S. Effects of different separation methods on the physical and functional properties of extracellular vesicles. PLOS ONE 2020;15:e0235793. [DOI: 10.1371/journal.pone.0235793].

29. Takov K, Yellon DM, Davidson SM. Comparison of small extracellular vesicles isolated from plasma by ultracentrifugation or size-exclusion chromatography: yield, purity and functional potential. Journal of Extracellular Vesicles 2019;8:1560809. [DOI: 10.1080/20013078.2018.1560809].

30. Radeghieri A, Alacqua S, Zendrini A, et al. Active antithrombin glycoforms are selectively physiosorbed on plasma extracellular vesicles. Journal of Extracellular Biology 2022;1:e57. [DOI: 10.1002/jex2.57].

31. T T, D B, S L, et al. Extracellular Vesicle Separation Techniques Impact Results from Human Blood Samples: Considerations for Diagnostic Applications. International journal of molecular sciences 2021;22. [PMID: 34502122 DOI: 10.3390/ijms22179211].

32. Williams C, Palviainen M, Reichardt N-C, Siljander PR-M, Falcón-Pérez JM. Metabolomics Applied to the Study of Extracellular Vesicles. Metabolites 2019;9:276. [DOI: 10.3390/metabo9110276].

33. Paolini L, Zendrini A, Noto GD, et al. Residual matrix from different separation techniques impacts exosome biological activity. Sci Rep 2016;6:23550. [DOI: 10.1038/srep23550].

34. Mol EA, Goumans M-J, Doevendans PA, Sluijter JPG, Vader P. Higher functionality of extracellular vesicles isolated using size-exclusion chromatography compared to ultracentrifugation. Nanomedicine: Nanotechnology, Biology and Medicine 2017;13:2061–5. [DOI: 10.1016/j.nano.2017.03.011].

35. Wolf M, Poupardin RW, Ebner-Peking P, et al. A functional corona around extracellular vesicles enhances angiogenesis, skin regeneration and immunomodulation. Journal of Extracellular Vesicles 2022;11:e12207. [DOI: 10.1002/jev2.12207].

36. Rome S. Muscle and Adipose Tissue Communicate with Extracellular Vesicles. Int J Mol Sci 2022;23:7052. [PMID: 35806052 DOI: 10.3390/ijms23137052].

37. Guescini M, Canonico B, Lucertini F, et al. Muscle Releases Alpha-Sarcoglycan Positive Extracellular Vesicles Carrying miRNAs in the Bloodstream. PLOS ONE 2015;10:e0125094. [DOI: 10.1371/journal.pone.0125094].

38. Estrada AL, Valenti ZJ, Hehn G, et al. Extracellular vesicle secretion is tissue-dependent ex vivo and skeletal muscle myofiber extracellular vesicles reach the circulation in vivo. American Journal of Physiology-Cell Physiology 2022;322:C246–59. [DOI: 10.1152/ajpcell.00580.2020].

39. Li Y, He X, Li Q, et al. EV-origin: Enumerating the tissue-cellular origin of circulating extracellular vesicles using exLR profile. Computational and Structural Biotechnology Journal 2020;18:2851–9. [DOI: 10.1016/j.csbj.2020.10.002].

40. Su Y, Chen M, Xu W, Gu P, Fan X. Advances in Extracellular-Vesicles-Based Diagnostic and Therapeutic Approaches for Ocular Diseases. ACS Nano 2024;18:22793–828. [DOI: 10.1021/acsnano.4c08486].

41. Mustapic M, Eitan E, Werner JK, et al. Plasma Extracellular Vesicles Enriched for Neuronal Origin: A Potential Window into Brain Pathologic Processes. Front Neurosci 2017;11. [DOI: 10.3389/fnins.2017.00278].

42. Latronico N, Rasulo FA, Eikermann M, Piva S. Publisher Correction to: Critical Illness Weakness, Polyneuropathy and Myopathy: Diagnosis, treatment, and longCterm outcomes. Crit Care 2023;27:469. [PMID: 38037109 DOI: 10.1186/s13054-023-04757-3].

43. Lad H, Saumur TM, Herridge MS, et al. Intensive Care Unit-Acquired Weakness: Not just Another Muscle Atrophying Condition. Int J Mol Sci 2020;21:7840. [PMID: 33105809 DOI: 10.3390/ijms21217840].

44. Faria M, Björnmalm M, Thurecht KJ, et al. Minimum information reporting in bio-nano experimental literature. Nat Nanotechnol 2018;13:777–85. [PMID: 30190620 DOI: 10.1038/s41565-018-0246-4].

45. Van Deun J, Mestdagh P, Agostinis P, et al. EV-TRACK: transparent reporting and centralizing knowledge in extracellular vesicle research. Nat Methods 2017;14:228–32. [DOI: 10.1038/nmeth.4185].

46. Lucien F, Gustafson D, Lenassi M, et al. MIBlood-EV: Minimal information to enhance the quality and reproducibility of blood extracellular vesicle research. Journal of Extracellular Vesicles 2023;12:12385. [DOI: 10.1002/jev2.12385].

47. Grossi I, Radeghieri A, Paolini L, et al. MicroRNAC34aC5p expression in the plasma and in its extracellular vesicle fractions in subjects with Parkinson’s disease: An exploratory study. Int J Mol Med 2021;47:533–46. [PMID: 33416118 DOI: 10.3892/ijmm.2020.4806].

48. Crescitelli R, Huang Y, Hendrix A, et al. Recommendations for Studying In Situ Extracellular Vesicles From Solid Tissue. Journal of Extracellular Vesicles 2025;14:e70185. [DOI: 10.1002/jev2.70185].

49. Paolini L, Radeghieri A, Civini S, Caimi L, Ricotta D. The epsilon hinge-ear region regulates membrane localization of the AP-4 complex. Traffic 2011;12:1604–19. [PMID: 21810154 DOI: 10.1111/j.1600-0854.2011.01262.x].

50. Salvi A, Vezzoli M, Busatto S, et al. Analysis of a nanoparticleCenriched fraction of plasma reveals miRNA candidates for Down syndrome pathogenesis. Int J Mol Med 2019;43:2303–18. [PMID: 31017260 DOI: 10.3892/ijmm.2019.4158].

51. Maiolo D, Paolini L, Di Noto G, et al. Colorimetric nanoplasmonic assay to determine purity and titrate extracellular vesicles. Anal Chem 2015;87:4168–76. [PMID: 25674701 DOI: 10.1021/ac504861d].

52. Zendrini A, Paolini L, Busatto S, et al. Augmented COlorimetric NANoplasmonic (CONAN) Method for Grading Purity and Determine Concentration of EV Microliter Volume Solutions. Frontiers in Bioengineering and Biotechnology 2020;7.

53. Ridolfi A, Brucale M, Montis C, et al. AFM-Based High-Throughput Nanomechanical Screening of Single Extracellular Vesicles. Anal Chem 2020;92:10274–82. [DOI: 10.1021/acs.analchem.9b05716].

54. Ridolfi A, Conti L, Brucale M, et al. Particle profiling of EV-lipoprotein mixtures by AFM nanomechanical imaging. Journal of Extracellular Vesicles 2023;12:12349. [DOI: 10.1002/jev2.12349].

55. Nečas D, Klapetek P. Gwyddion: an open-source software for SPM data analysis. Open Physics 2012;10:181–8. [DOI: 10.2478/s11534-011-0096-2].

56. Busatto S, Zendrini A, Radeghieri A, et al. The nanostructured secretome. Biomater Sci 2019;8:39–63. [DOI: 10.1039/C9BM01007F].

57. Zhang H, Freitas D, Kim HS, et al. Identification of distinct nanoparticles and subsets of extracellular vesicles by asymmetric flow field-flow fractionation. Nat Cell Biol 2018;20:332–43. [DOI: 10.1038/s41556-018-0040-4].

58. Noto GD, Paolini L, Zendrini A, Radeghieri A, Caimi L, Ricotta D. C-src Enriched Serum Microvesicles Are Generated in Malignant Plasma Cell Dyscrasia. PLOS ONE 2013;8:e70811. [DOI: 10.1371/journal.pone.0070811].

59. Di Noto G, Chiarini M, Paolini L, et al. Immunoglobulin free light chains and GAGs mediate Multiple Myeloma Extracellular Vesicles uptake and secondary NfkB nuclear translocation. Frontiers in Immunology 2014;5.

60. Jeppesen DK, Zhang Q, Franklin JL, Coffey RJ. Extracellular vesicles and nanoparticles: emerging complexities. Trends in Cell Biology 2023;33:667–81. [DOI: 10.1016/j.tcb.2023.01.002].

61. Yamamoto T, Yamamoto A, Watanabe M, et al. Classification of FABP isoforms and tissues based on quantitative evaluation of transcript levels of these isoforms in various rat tissues. Biotechnol Lett 2009;31:1695–701. [DOI: 10.1007/s10529-009-0065-7].

62. Paolini L, Orizio F, Busatto S, et al. Exosomes Secreted by HeLa Cells Shuttle on Their Surface the Plasma Membrane-Associated Sialidase NEU3. Biochemistry 2017;56:6401–8. [DOI: 10.1021/acs.biochem.7b00665].

63. Kowal J, Arras G, Colombo M, et al. Proteomic comparison defines novel markers to characterize heterogeneous populations of extracellular vesicle subtypes. Proceedings of the National Academy of Sciences 2016;113:E968–77. [DOI: 10.1073/pnas.1521230113].

64. Lötvall J, Hill AF, Hochberg F, et al. Minimal experimental requirements for definition of extracellular vesicles and their functions: a position statement from the International Society for Extracellular Vesicles. Journal of Extracellular Vesicles 2014;3:26913. [PMID: 25536934 DOI: 10.3402/jev.v3.26913].

65. Musicò A, Zenatelli R, Romano M, et al. Surface functionalization of extracellular vesicle nanoparticles with antibodies: a first study on the protein corona “variable.” Nanoscale Advances 2023;5:4703–17. [DOI: 10.1039/D3NA00280B].

66. Paolini L, Federici S, Consoli G, et al. Fourier-transform Infrared (FT-IR) spectroscopy fingerprints subpopulations of extracellular vesicles of different sizes and cellular origin. Journal of Extracellular Vesicles 2020;9:1741174. [DOI: 10.1080/20013078.2020.1741174].

67. Mangolini V, Gualerzi A, Picciolini S, et al. Biochemical Characterization of Human Salivary Extracellular Vesicles as a Valuable Source of Biomarkers. Biology 2023;12:227. [DOI: 10.3390/biology12020227].

68. Huang Y, Driedonks TAP, Cheng L, et al. Brain Tissue-Derived Extracellular Vesicles in Alzheimer’s Disease Display Altered Key Protein Levels Including Cell Type-Specific Markers. J Alzheimers Dis 2022;90:1057–72. [PMID: 36213994 DOI: 10.3233/JAD-220322].

69. Borup A, Boysen AT, Ridolfi A, et al. Comparison of separation methods for immunomodulatory extracellular vesicles from helminths. Journal of Extracellular Biology 2022;1:e41. [DOI: 10.1002/jex2.41].

70. Montis C, Zendrini A, Valle F, et al. Size distribution of extracellular vesicles by optical correlation techniques. Colloids and Surfaces B: Biointerfaces 2017;158:331–8. [DOI: 10.1016/j.colsurfb.2017.06.047].

71. Busatto S, Yang Y, Walker SA, et al. Brain metastases-derived extracellular vesicles induce binding and aggregation of low-density lipoprotein. Journal of Nanobiotechnology 2020;18:162. [DOI: 10.1186/s12951-020-00722-2].

72. Rogers NMK, McCumber AW, McMillan HM, et al. Comparative electrokinetic properties of extracellular vesicles produced by yeast and bacteria. Colloids and Surfaces B: Biointerfaces 2023;225:113249. [DOI: 10.1016/j.colsurfb.2023.113249].

73. Musicò A, Zendrini A, Reyes SG, et al. Extracellular vesicles of different cellular origin feature distinct biomolecular corona dynamics. Nanoscale Horiz 2024;10:104–12. [DOI: 10.1039/D4NH00320A].

74. Radeghieri A, Bergese P. The biomolecular corona of extracellular nanoparticles holds new promises for advancing clinical molecular diagnostics. Expert Rev Mol Diagn 2023;23:471–4. [PMID: 37199178 DOI: 10.1080/14737159.2023.2215927].

75. Paolini L, Zendrini A, Radeghieri A. Biophysical properties of extracellular vesicles in diagnostics. Biomark Med 2018;12:383–91. [PMID: 29441794 DOI: 10.2217/bmm-2017-0458].

76. Huang Y, Cheng L, Turchinovich A, et al. Influence of species and processing parameters on recovery and content of brain tissue-derived extracellular vesicles. J Extracell Vesicles 9:1785746. [PMID: 32944174 DOI: 10.1080/20013078.2020.1785746].

77. Moore HM, Kelly A, Jewell SD, et al. Biospecimen Reporting for Improved Study Quality. Biopreserv Biobank 2011;9:57–70. [PMID: 21826252 DOI: 10.1089/bio.2010.0036].

78. Mucci NR, Moore HM, Brigham LE, et al. Meeting research needs with postmortem biospecimen donation: summary of recommendations for postmortem recovery of normal human biospecimens for research. Biopreserv Biobank 2013;11:77–82. [PMID: 24845428 DOI: 10.1089/bio.2012.0063].

79. Shen S, Shen Z, Wang C, et al. Effects of lysate/tissue storage at −80°C on subsequently extracted EVs of epithelial ovarian cancer tissue origins. iScience 2023;26:106521. [DOI: 10.1016/j.isci.2023.106521].

80. D’Arrigo D, Lässer C, Urzì O, et al. Extracellular vesicles isolated from frozen and fresh human melanoma tissue are similar in purity and protein composition. 2024:2024.04.03.587936.

