## Supplementary files for "Shared protocol to enrich and compare biochemical and biophysical properties of Extracellular Vesicles from human plasma and skeletal muscle biopsy"

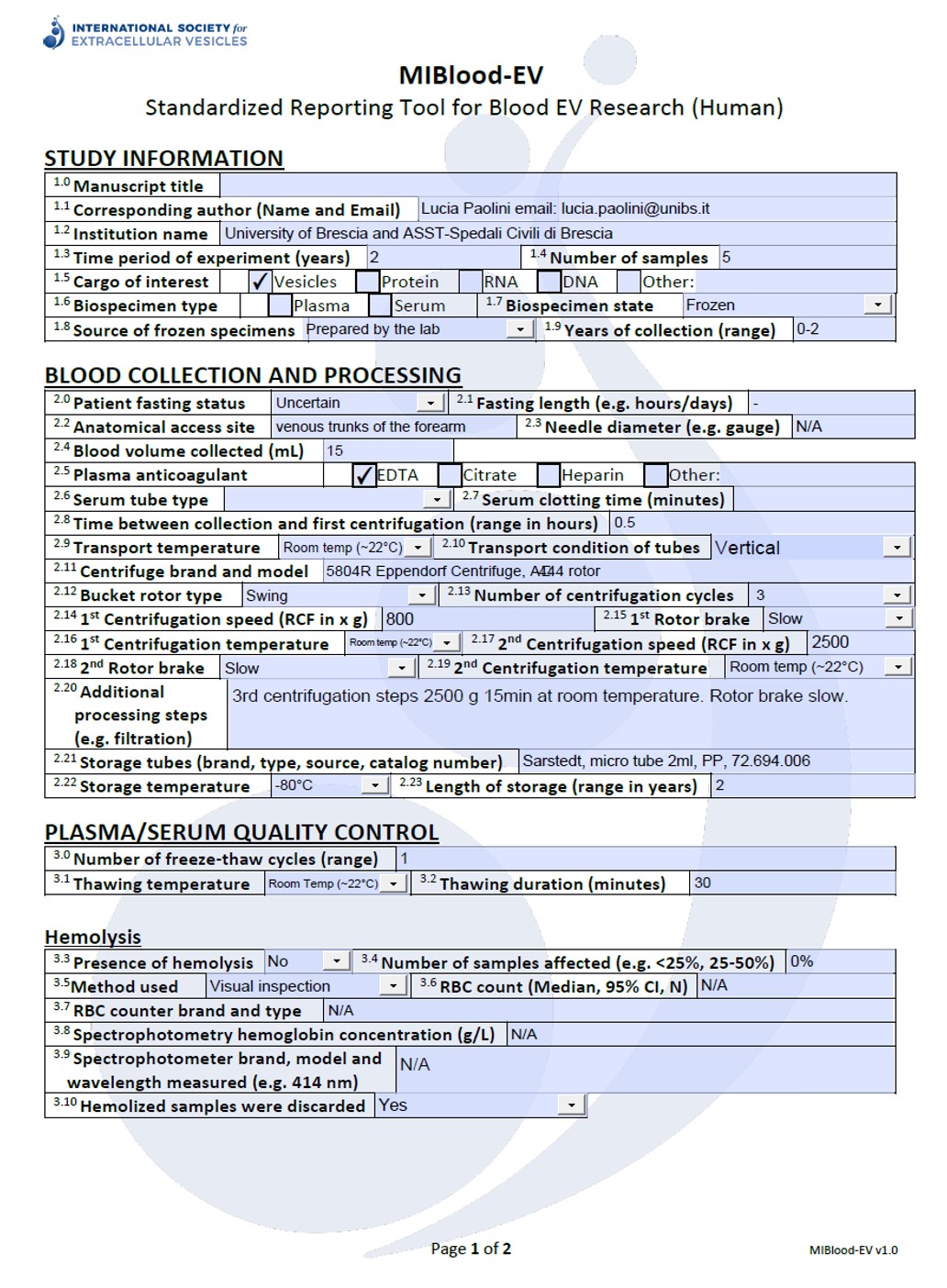

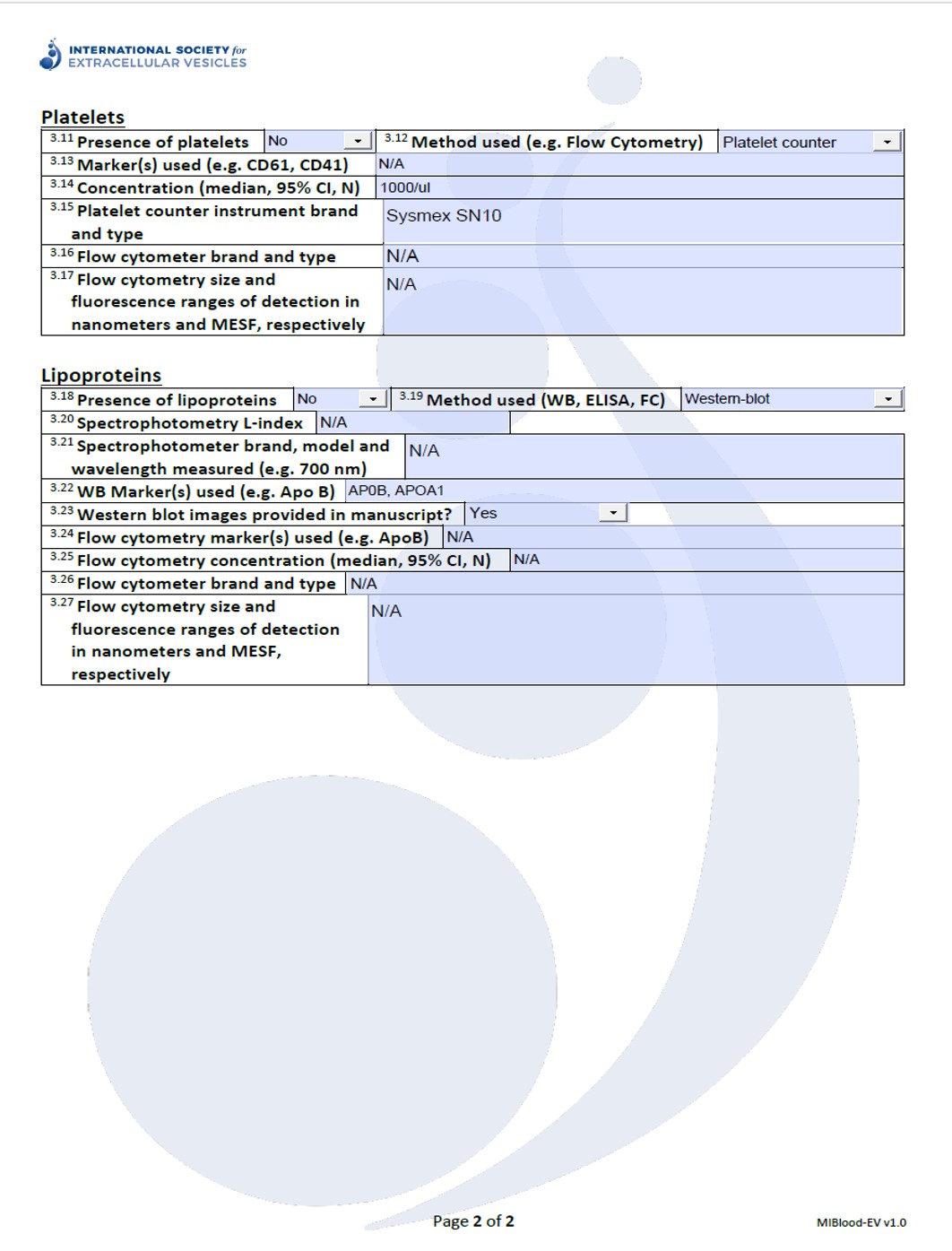

Fig.1SI: MIBlood survey. Survey completed according to Minimal information to enhance the quality and reproducibility of blood extracellular vesicle research^[1]^.

**Supplementary Methods**

**Subject Recruitment and Sample Collection**

In this study, 19 plasma samples and 12 skeletal muscle (SkM) tissue biopsies were obtained from subjects hospitalized in the Intensive Care Unit (n=14) and Orthopedic ward (n=5) of the Spedali Civili University Hospital in Brescia, Italy. The Ethics Committee of Brescia approved the study in adherence with the Declaration of Helsinki and Good Clinical Practice (Protocol number 34 68). All traceable identifiers were removed before analysis to protect patient confidentiality; all samples were analyzed anonymously. Written informed consent was obtained from the patient before the collection of samples. In case of altered consciousness, the Ethics Committees waived the requirement for consent, as in Italy relatives are not regarded as legal representatives of the patient in the absence of a formal designation. Written informed consent was requested from all surviving patients as soon as they regained their mental competency. The study cohort consisted of 19 subjects, including 14 males (73.7%) and 5 females (26.3%), with an age range of 38 – 83 years. (average age 68,2 years). The mean age was 67 years in males and 71,6 years in females. Both blood and SkM tissue biopsies were collected at the same time and processed within 30 minutes from the withdrawal.

Plasma was obtained as previously described from peripheral blood samples in EDTA^[2]^. Blood was collected through a peripheral venous catheter in one of the main venous trunks of the forearm. Careful tube transportation was ensured to avoid unnecessary agitation and blood was kept at room temperature. Briefly blood was centrifuged at 800 x g for 10 min (5804R Eppendorf Centrifuge, A‑4‑44 rotor, 15 ml tubes), 2,500 x g for 15 min and then centrifuged a second time at 2,500 x g for 15 min to obtain platelets and cell-depleted plasma for further analyses^[2–5]^. All centrifugations steps were made at room temperature with 4 of acceleration and 3 of brake. After each centrifugation plasma was collected in a fresh plastic tube, leaving 1 cm of plasma above the buffy layer so as not to disturb it. Plasma was finally transferred into crio-vials in 1 ml aliquots, snap-frozen in liquid nitrogen and stored at ‑80˚C.

Human SkM tissue was dissected from quadriceps femoris. The muscle samples harvested from patients hospitalized in ICU were obtained with a surgical biopsy. The skin was prepped with antiseptic Clorhexidin 2% solution and draped. A longitudinal 4 cm midline incision was performed at the middle third of the thigh. The quadriceps muscular fascia was reached through the subcutaneous tissue and longitudinally incised. A 2x1x1 cm sample of rectus femoris muscular tissue was harvested with a n° 21 blade. Hemostasis was obtained in every case by compression. The quadriceps fascia and the subcutaneous tissue were sutured with 3-0 polyglactin 910 braided adsorbable suture; the skin was sutured with 4-0 polyglactin 910 braided adsorbable suture. A compressive dressing was applied at the end of the procedure. The samples obtained from patients in the Orthopeadic Ward were obtained during total hip replacement performed for hip arthritis through a direct anterior approach. Fibres of the rectus femoris were harvested during the exposure of the hip joint, in the deep portion of the interval between sartorius and tensor fasciae latae. A 2x1x1 cm sample was obtained with a n° 21 scalpel. As soon as the biopsy was removed from the patient, it was immediately placed into a polypropylene tube filled with 25 ml cold (+4°C) NaCl 0.9% solution and washed twice. SkM tissue was, then, moved to a new tube and snap-frozen in liquid nitrogen. Samples was stored at -80°C.

**Enrichment of EV from human plasma**

Plasma EVs were separated through serial ultra-centrifugation and discontinuous sucrose gradient (DSG). One ml aliquot was processed, in parallel, with serial centrifugation steps. Briefly, 1 ml plasma was centrifuged at 300 x g for 10 min (5417C Eppendorf Centrifuge, 45‑30‑11 rotor, 1.5 ml Eppendorf tubes, 1 ml each tube). Supernatant (1 ml) was transferred to a new tube and centrifuged at 2,000 x g for 20 min (5417C Eppendorf Centrifuge, 45‑30‑11 rotor, 1.5 ml Eppendorf tubes, 1 ml each tube). Supernatant (1 ml) was transferred to a new tube and centrifuged at 16,500 x g for 20 min (5417C Eppendorf Centrifuge, 45‑30‑11 rotor, 1.5 ml Eppendorf tubes, 1 ml each tube) and finally supernatant was transferred to an appropriate tube and centrifuged at 118,000 x g for 2h and 30 min (Optima MAX, TLA‑55 rotor, 1.5 ml polypropylene microfuge tube, Beckman). The 16,500 x g centrifugation step allows sedimentation of 16k sample, while the 118,000 x g ultracentrifugation step enriches 118k sample.

DSG was carried out as follows: Plasma 16k and Plasma 118k pellet, obtained as described above, were further processed adapting the protocols developed in previous studies^[2,6]^. Briefly, 16k and 118k were re‑suspended, separately, in 1ml buffer A (10 mM Tris‑HCl 250 mM sucrose, pH 7.4), loaded on top of a discontinuous sucrose gradient 15% (600 μl), 20, 25, 30, 40, 60, 65% (400 μl), 70% (800 μl) sucrose in 10 mM Tris‑HCl, pH 7.4) and centrifuged at 230,000 x g for 16 h at 4˚C (rotor MLS 50; Beckman Optima MAX, no brake). Twelve fractions with equal volumes (400 μl) were collected from the top of the gradient. All fractions were diluted with 600 μl water suitable for High-performance liquid chromatography (HPLC water). For Western blot analysis fractions 1-12 (250 μl) were precipitated by incubation with 10% trichloracetic acid (TCA) (Sigma) for 4 h as described in Paolini et al^.[7]^. For all the other analyses, fractions from 6 to 9 (750 μl each) were ultracentrifuged (100,000 x g for 2 h at 4˚C, Optima MAX, TLA‑55 rotor, 1.5 ml polypropylene microfuge tube, Beckman) and resupended in 100 μl HPLC water (total volume; HPLC water was previously ultracentrifuged at 100,000 g for 2 h in Optima MAX, TLA‑55 rotor, 1.5 ml polypropylene microfuge tube, Beckman) and aliquoted for further analyses.

**Enrichment of EVs from human SkM tissue**

EVs were separated from human SkM tissue after mechanical and enzymatic dissociation, as previously described from Crescitelli et al.^[8]^. with some modifications. Briefly, frozen tissue was thawed 2 min at room temperature, washed one with cold PBS1x and weighted (5 samples, mean 0.89 gr). Tissue was transferred in a 100 mm dish with cold RPMI (medium/tissue ratio: 1ml RPMI/ 0.2 gr tissue) and gently cut into small pieces (2x2x2mm) with a scalpel. Tissue pieces were transferred in a 6-well plate with 0.2 gr of SkM tissue each well. RPMI medium was added to each well to reach 2 ml of total medium. SkM tissue was incubated with DNase (Roche, 40U/ml final concentration, catalogue number 11284932001, batch not available) and collagenase D (Roche, 2mg/ml final concentration. Catalogue number 11088858001, batch 66505623) at 37°C for 30 minutes in gently rotation (78 rpm). No enzyme inhibitors were added at this step to stop the reaction. This in order to not increase the concentration of soluble protein in the samples. Enzymatic reaction was inhibited diluting the samples with 3 times the starting volume of RPMI medium. Medium derived after the enzymatic dissociation was filtered in a 0,70 µm pore strain (Corning, Nylon, sterile, catalogue number 431751) in order to eliminate the largest debris. The filtrate was used for EVs isolation through serial ultra-centrifigation and DSG as described for plasma.

Briefly, Tissue filtrate was centrifuged at 300 x g for 10 min (5804R Eppendorf centrifuge,

A-4-44 rotor, 50 mL 174 × 22 mm polypropylene tube Sarstedt). Supernatant was transferred to a new tube and centrifuged at 2,000 x g for 20 min (5804R Eppendorf centrifuge, A-4-44 rotor, 50 mL 174 × 22 mm polypropylene tube Sarstedt). Again, supernatant was transferred to a new tube and centrifuged at 16,500 x g for 20 min in order to obtain Tissue 16K preparation (Avanti J25, rotor ja20, polycarbonate tubes 357003 Beckman). Finally, supernatant was transferred to an appropriate tube and ultracentrifuged at 118,000 x g for 2 h and 30 min in order to get Tissue 118K preparation (Optima XP80, TY45i rotor, polycarbonate tubes 355622 Beckmann). All centrifuges were performed at 4°C.

DSG was carried out as follows: Tissue 16k and Tissue 118k pellet obtained were treated as described for plasma preparations.

For Western blot analysis fractions 1-12 (250 μl) were precipitated by incubation with 10% trichloracetic acid (TCA) (Sigma) for 4 h as described in Paolini et al.^7^. For all the other analyses, fractions from 6 to 9 (750 μl each) were ultracentrifuged (100,000 x g for 2 h at 4˚C, Optima MAX, TLA‑55 rotor, 1.5 ml polypropylene microfuge tube, Beckman) and resupended in 100 μl HPLC water (total volume; HPLC water was previously ultracentrifuged at 100,000 g for 2 h in Optima MAX, TLA‑55 rotor, 1.5 ml polypropylene microfuge tube, Beckman) and aliquoted for further analyses.

SkM tissue homogenate (TH) was obtained from SkM tissue after the mechanical and enzymatic dissociation. Briefly, SkM was resuspended in cold RIPA buffer 1X + Protease Inibitors (500 ul each 0.1 gr of tissue) and homogenized with Ultra-Turrax T25 basic (IKA-Werke) homogenizer for 1 minute in ice. Sample was centrifuged at 800 x g 15 min at 4°C twice. Supernatant was collected and kept for protein quantification and SDS-PAGE analyses. In alternative, SkM homogenate was obtained with the same protocol, from tissue before the mechanical and enzymatic dissociation (THB).

**SDS-PAGE and Western Blot analysis**

SDS−PAGE and Western blot were performed by standard procedures^[9,10]^ on total plasma or tissue homogenate (30 μg, protein content determined by Bradford assay) and sucrose density gradient fractions from 1 to 12 (equal volumes of each fraction were loaded on a acrylamide–bisacrylamide gel). Samples were electrophoresed on Acrylamide-bisacrylamide gels 12.5% and analysed by Western Blot with the following antibodies^[11,12]^ (at dilution 1:500): mouse anti-Flotillin 1 (Santa Cruz, clone C-2, sc-74566), mouse anti-CD63 (Millipore, clone RFAC4, CBL553), rabbit anti-ADAM10 (Origene, AP05830PU-N), mouse anti-CD81 (Santa Cruz Biotechnology, clone B11, sc-166029), Caveolin 3 (Santa Cruz Biotechnology, sc-5310), Beta enolase (Santa Cruz Biotechnology, sc-100811), TSG101 (Santa Cruz Biotechnology, clone C‑2, sc‑7964), APOB (Santa Cruz Biotechnology, sc-393636) APOA1 (Thermo Scientific Fisher, 701239), FABP (Santa Cruz Biotechnology, sc-271529) Mouse anti-GM130 Cis-Golgi protein (BD Transduction, clone 35/130, 610822), Mouse anti-Hemoglobin A (Abnova, clone 4F9) were diluted 1:1000.

**Colorimetric nanoplasmonic (CONAN) assay**

The EV preparations (fraction 6-9 of the sucrose gradient) were checked for purity from protein contaminants using the COlorimetric NANoplasmonic (CONAN) assay, a recognized method to determine the purity of EV preparations^[13]^ that is based on the clustering of gold NPs onto lipid membranes^[14]^. The CONAN assay was performed as previously described in Zendrini et al.^[15]^. Briefly, EVs were resuspended in 100 μl of HPLC water. Two microlitre of EV solution at serial dilutions in HPLC water (1:1, 2μl of starting sample – 1:10, 2 μl of sample diluted in HPLC H_2_O – 1:100, 2 μl of sample diluted in HPLC H_2_O) were resuspended in 23 μl of water, mixed with 50 μl of AuNPs 6 nM and 25 μl of PBS. The result of the assay was collected on Ensight Multi Mode Reader (Perkin Elmer). Measurements were performed for each sample in triplicate.

**Bicinchoninic acid (BCA) and Bradford assay**

Protein concentrations of samples (fractions 6-9) were determined with a Pierce™ BCA Protein Assay Kit (ThermoFisher, Rockford, USA), and Bradford assay (Biorad) following the manufacturer's instructions.

**Atomic Force Microscopy (AFM) analysis**

AFM imaging was used to determine sample morphology and quantitative AFM morphometry was performed as described elsewhere^[16,17]^. Briefly, 5 μL aliquots from samples were deposited on glass coverslips previously functionalized with 0.01 mg/mL poly-L-lysine and left to adsorb for 60’ at 4°C. Aliquots were progressively diluted up to 200x in successive depositions in order to maximise the surface density of isolated particles. Imaging was performed in ultrapure water at room temperature on a Bruker Multimode8 AFM equipped with a Nanoscope V controller, a sealed fluid cell and a type JV piezoelectric scanner using Bruker ScanAsystFluid+ probes. Background subtraction was performed using Gwyddion^[18]^ 2.61. Quantitative morphometry was performed with custom Python scripts to recover the surface contact angle (CA) and equivalent solution diameter (D) of several hundred individual objects for each sample. The relative particle concentrations of different samples were estimated as follows. For each AFM image, particles were counted and divided by the scanned area to recover the image’s particle surface density in particles per square micron. This figure was then multiplied by the dilution factor employed for the deposition (see above) to recover a concentration factor which we demonstrated to be proportional to the original concentration of the deposited sample^[19]^, allowing us to quantitatively estimate the relative particle concentration across several samples.

**Nanotracking particle analysis (NTA)**

Pellets from 6 to 9 obtained from both plasma and human SkM tissue were resuspended in 100 µl of sterile HPLC water (Milli-Q; Meck Millipore) and pooled together for the characterization by Nanoparticles Tracking Analysis (NTA) using a NanoSight NS300 system (Malvern, Panalytical LTD, Malvern, UK) to evaluate the concentration and size distribution of EVs. The system was equipped with a Blue488 laser and sCMOS camera. Before samples analysis, filtered PBS (0,22µm filtered) was analysed for particle contamination. For the analysis, all samples were diluted in filtered PBS (plasma EVs 1:200 – human SkM EVs 1:500) to a final volume of 1 ml to obtain an optimal particles per frame value (20-120 particles/frame). EVs were injected in the sample chamber through a Nanosight syringe pump (Malvern, Panalytical LTD, Malvern, UK) that provides a continuous flow (50µl/min) using a 1 ml syringe at room temperature. Recordings of the movements of particles were collected for 60 s.

**Dynamic light scattering (DLS) analysis and zeta potential measurement**

DLS at θ = 90° and zeta potential measurements were performed using a Brookhaven Instrument 90 Plus (Brookhaven Instruments Corporation, Holtsville, NY). For DLS, each measurement was an average of four replicates of 1 minute each. The autocorrelation functions were analysed by Laplace inversion using the CONTIN algorithm, allowing an estimate of the particles size distribution. Zeta potentials were obtained from the electrophoretic mobility u, according to the Helmholtz-Smoluchowski equation: ζ = (η⁄ε) × u where η is the viscosity of the medium and ε is the dielectric permittivity of the dispersing medium. The zeta potential values are given as the average of ten measurements.

**Supplementary results**

**Biochemical Characterization**

Western Blot analyses was performed as indicated in methods section above. Un-cropped version of WB showed in main text (Fig. 2 and Fig. 3) are presented in this section. Biochemical characterization of Gradient fractions of Plasma 16k and 118k samples are showed in Fig 2-9SI; Gradient fractions of Tissue 16k and 118k samples are showed in Fig.10-15SI. Antibodies and dilutions as indicated in the figure or in SI methods section.

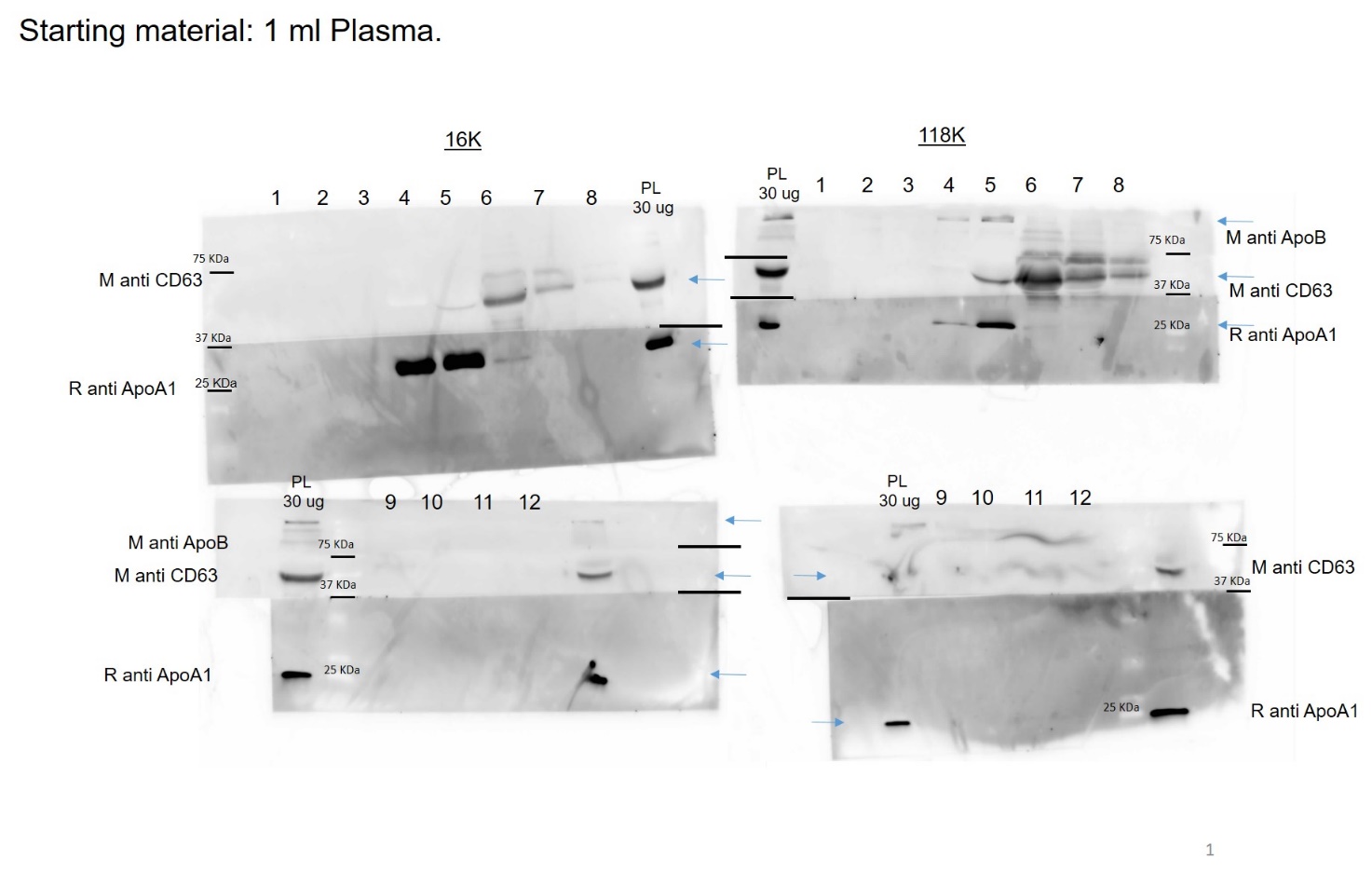

Fig.2SI. WB of Plasma 16k and 118k sample. 1-12 SDG fractions; PL: total plasma

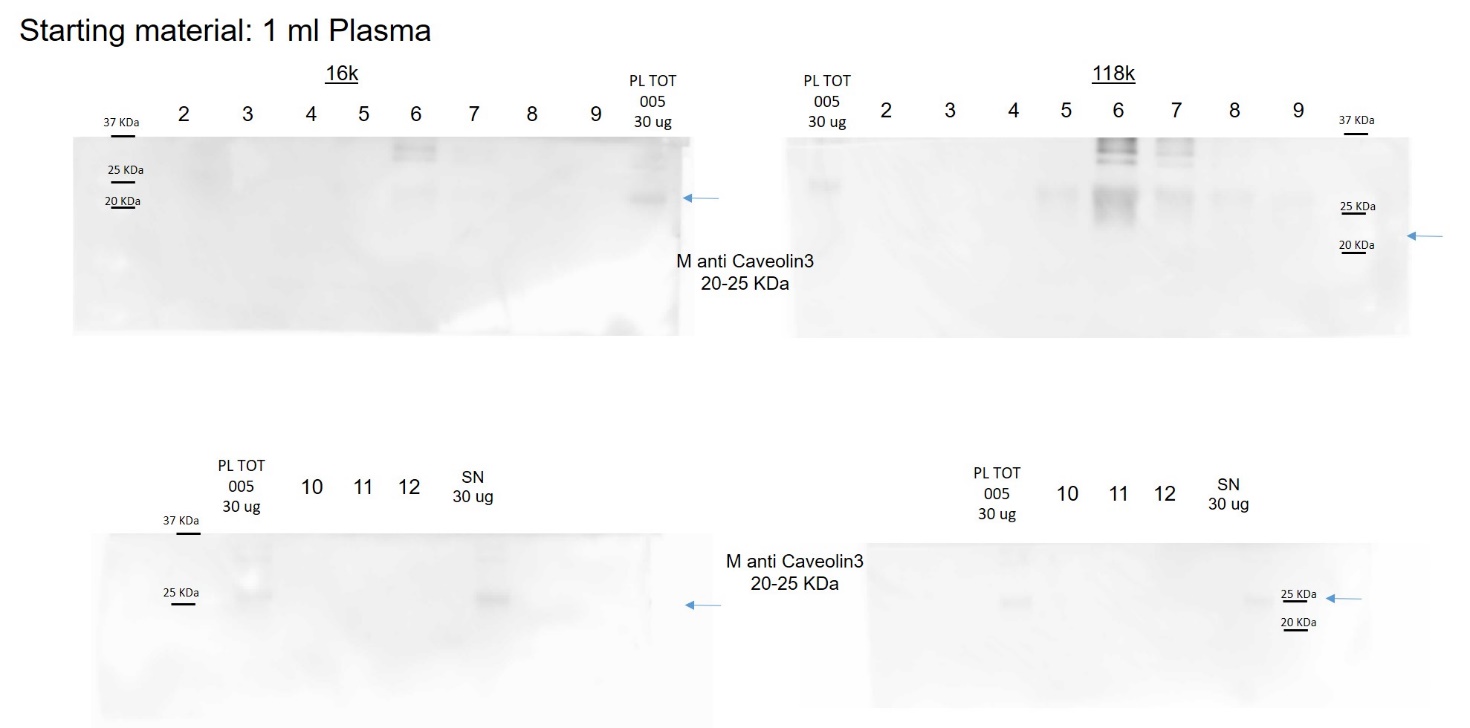

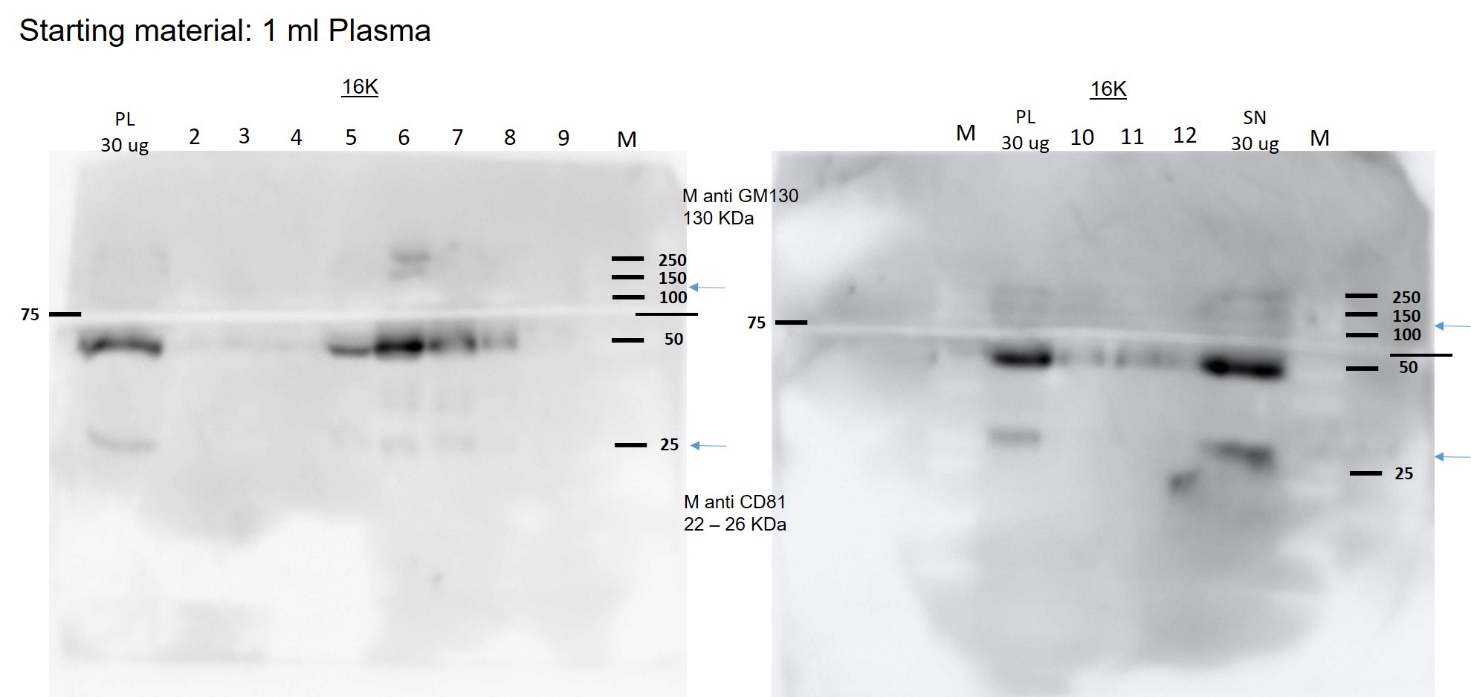

Fig.3SI. WB of Plasma 16k and 118k sample. 1-12 SDG fractions; PL: total plasma

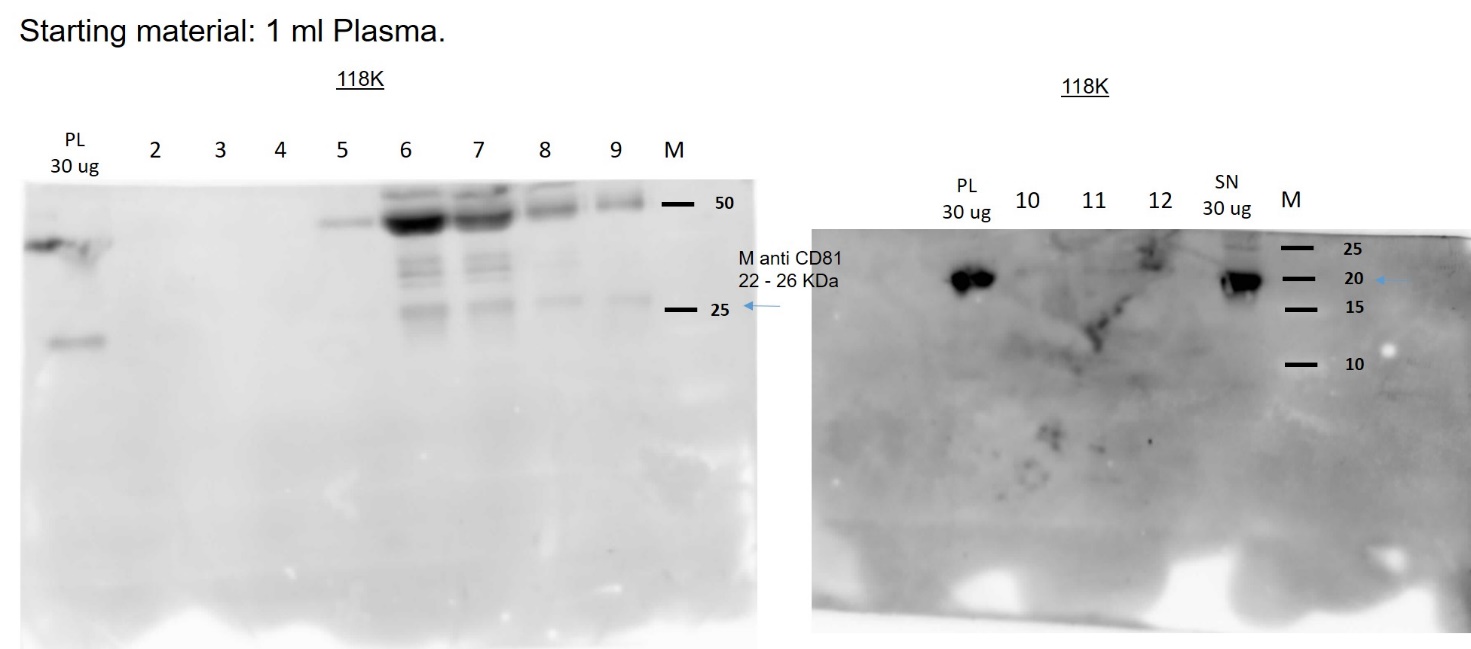

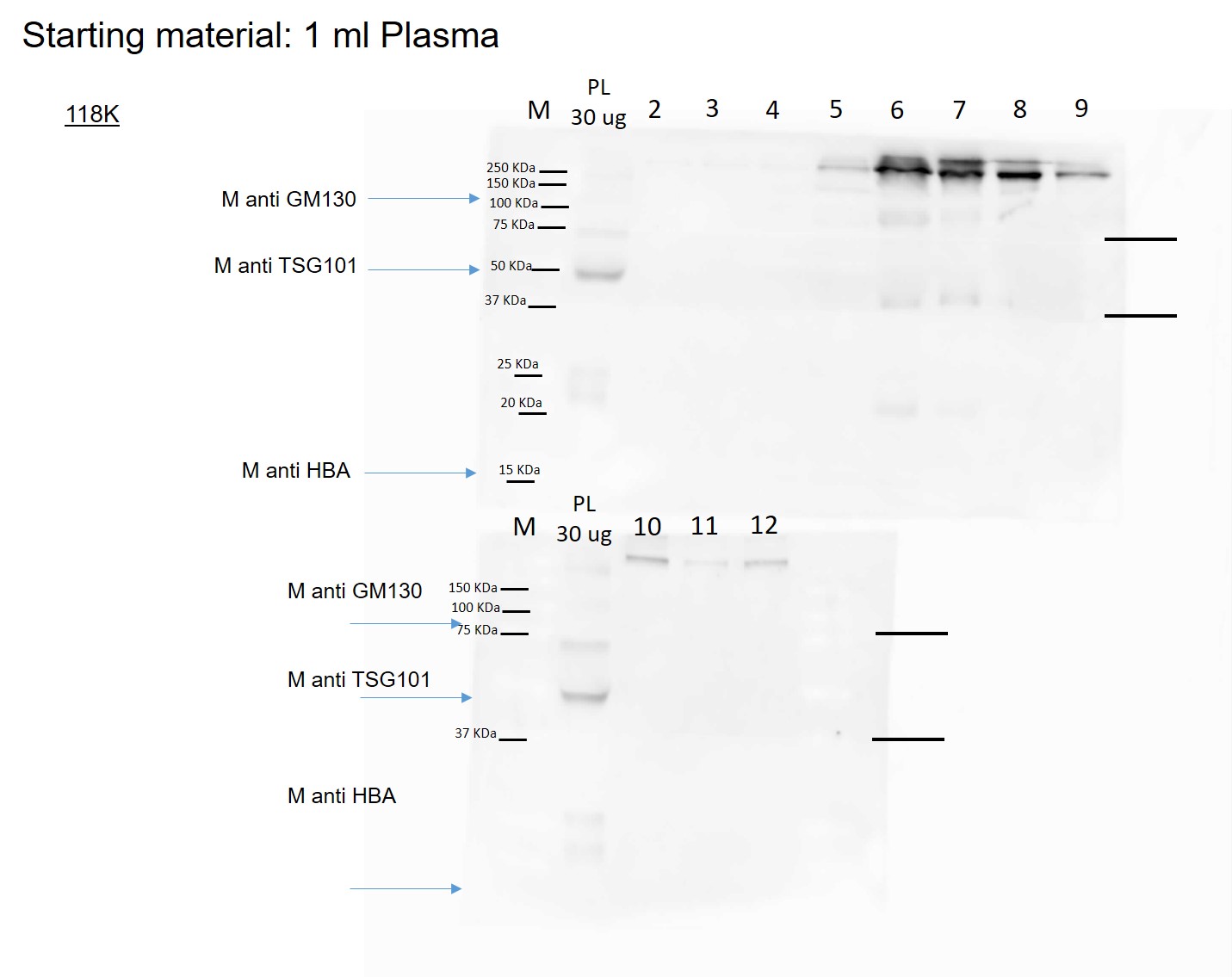

Fig.4SI. WB of Plasma 16k and 118k sample. 1-12 SDG fractions; PL: total plasma.

HBA: Hemoglobin A.

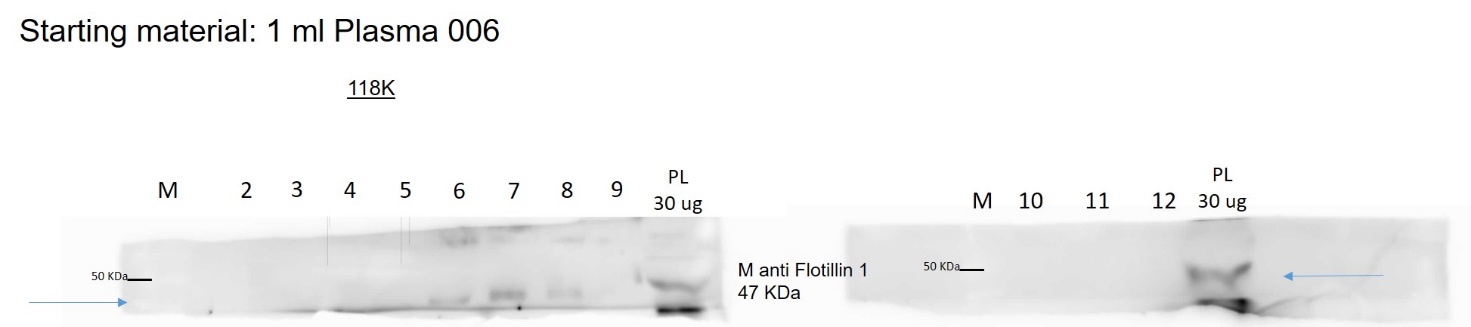

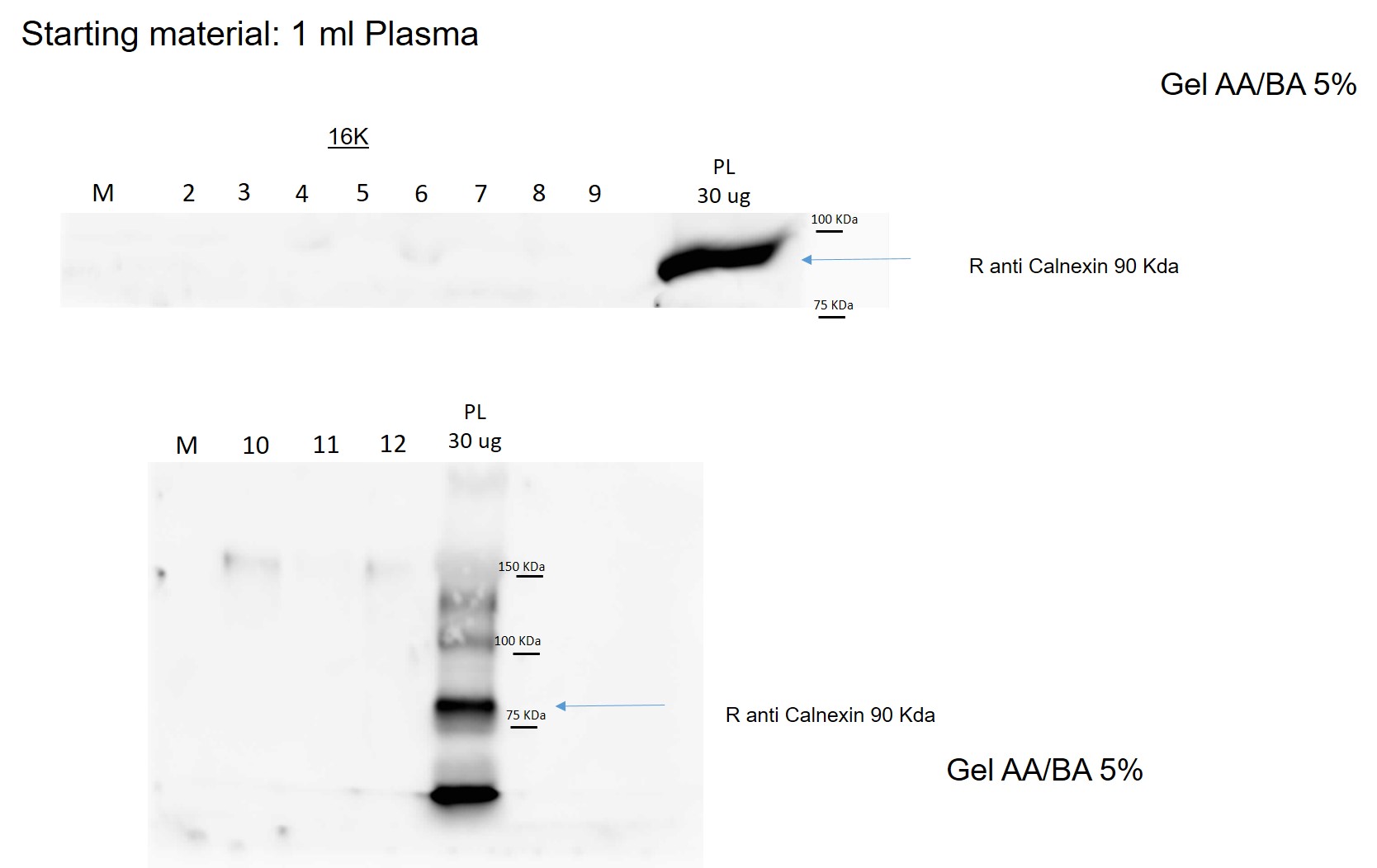

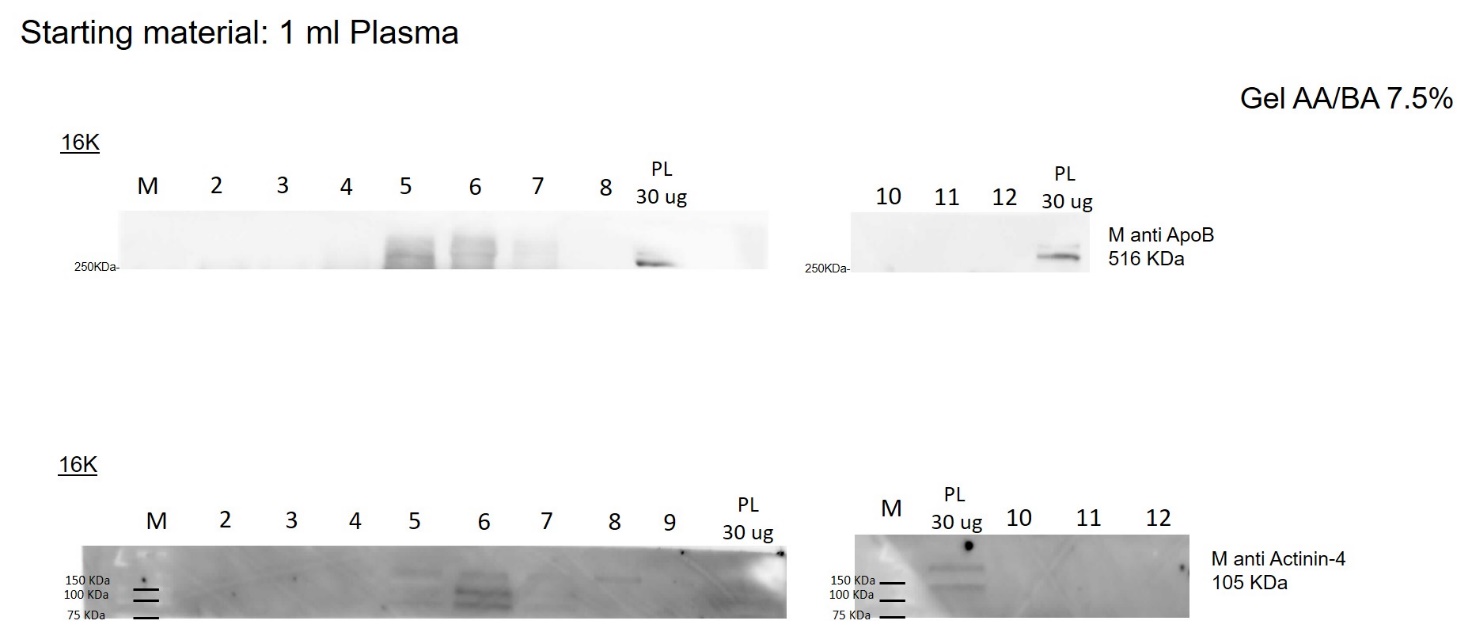

Fig.5SI. WB of Plasma 16k and 118k sample. 1-12 SDG fractions; PL: total plasma.

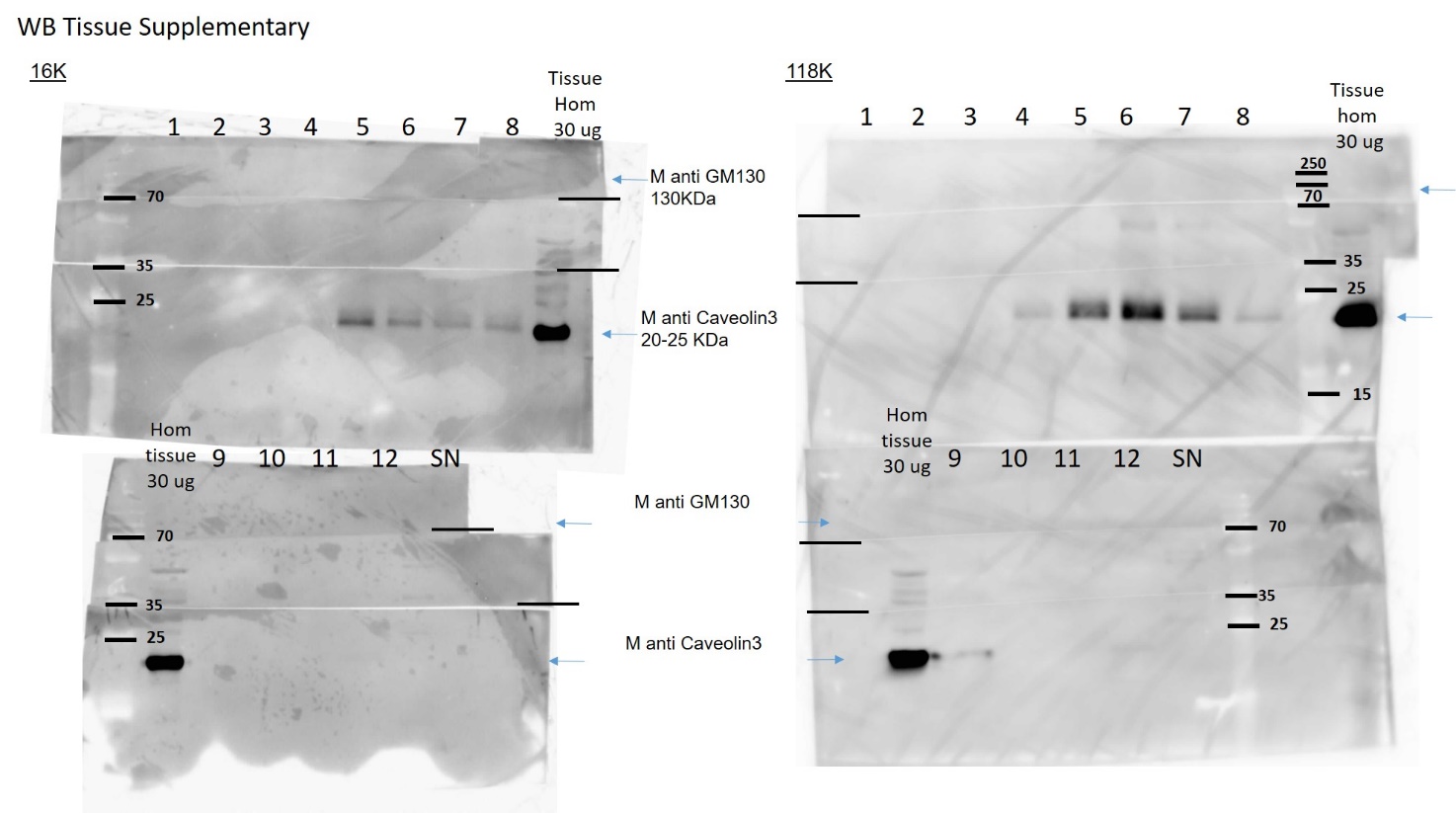

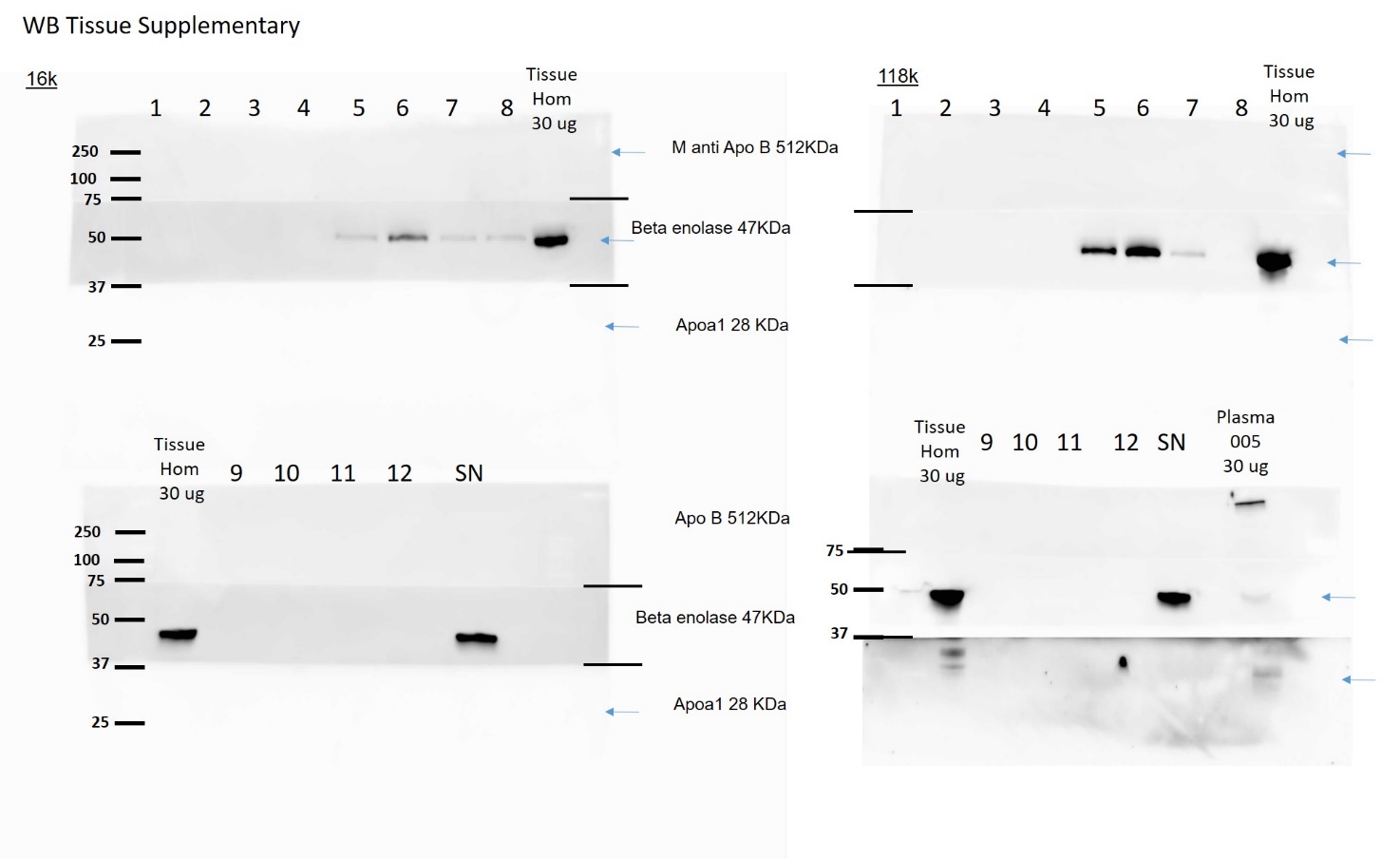

Fig.6SI. WB of Tissue 16k and 118k sample after sucrose density gradient (SDG). 1-12 SDG fractions; Tissue Hom: Skeletal muscle tissue homogenate after enzymatic and mechanical treatment; PL: total plasma

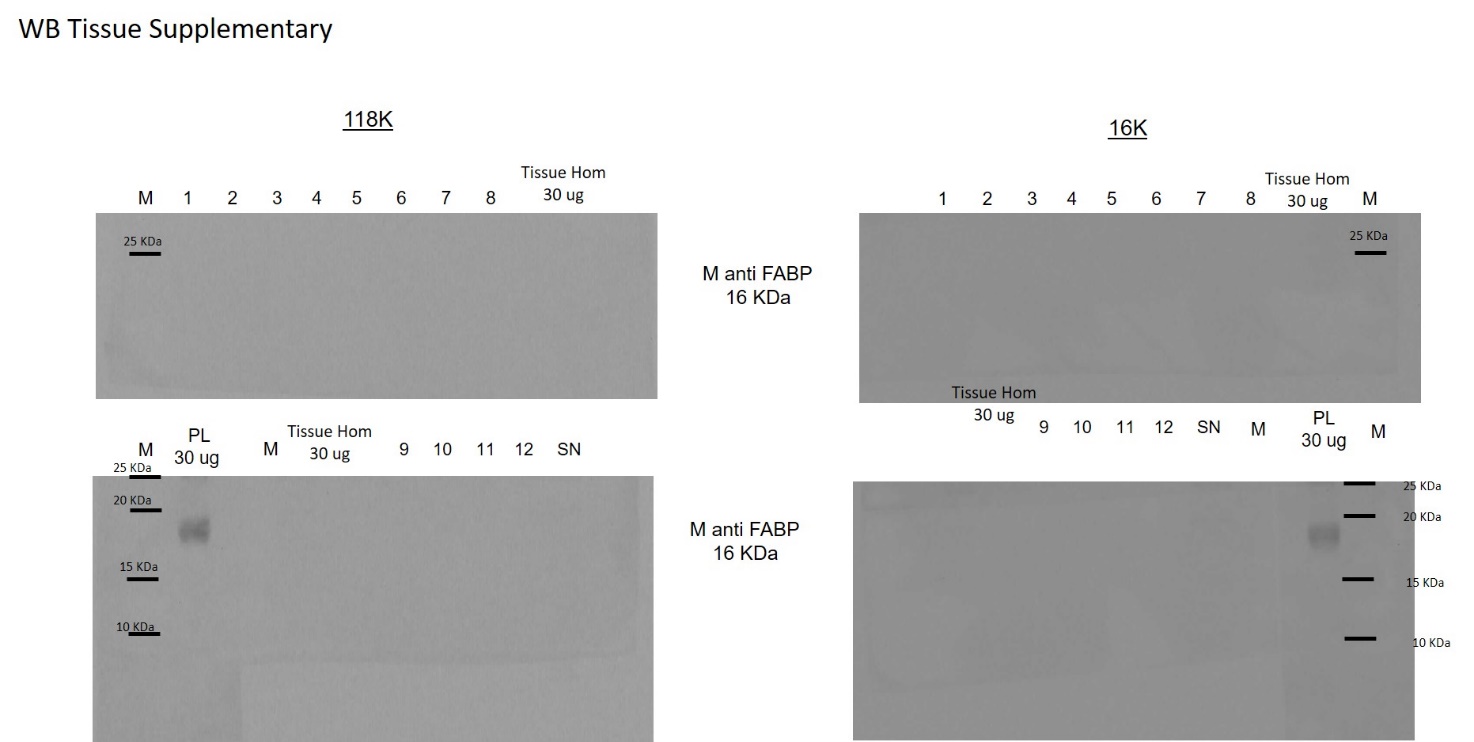

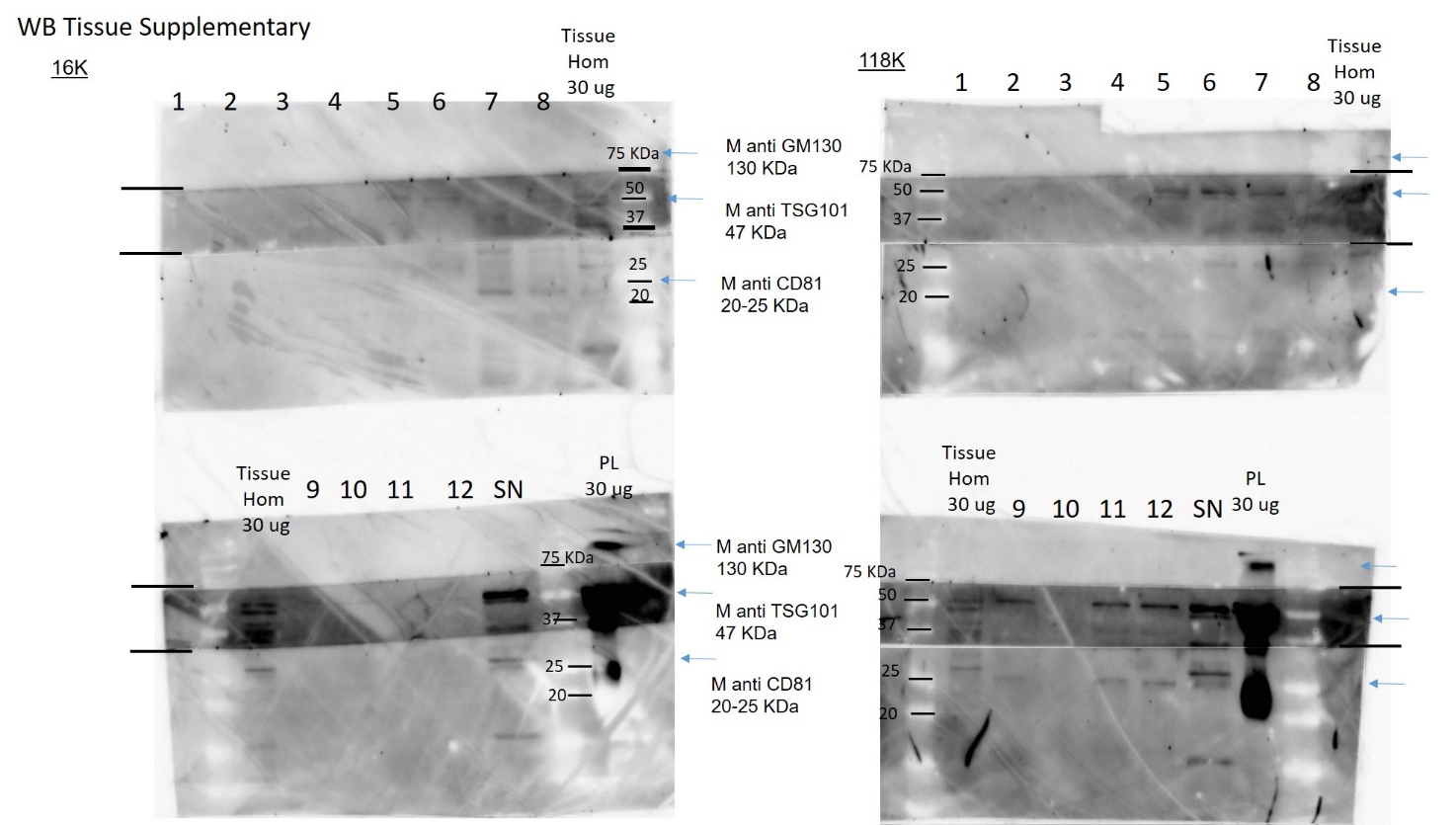

Fig.7SI. WB of Tissue 16k and 118k sample. DSG 1-12 sucrose fractions. PL: total plasma; Tissue Hom: Skeletal muscle tissue homogenate after mechanical and enzymatic dissociation; FABP: Fatty Acid Binding protein

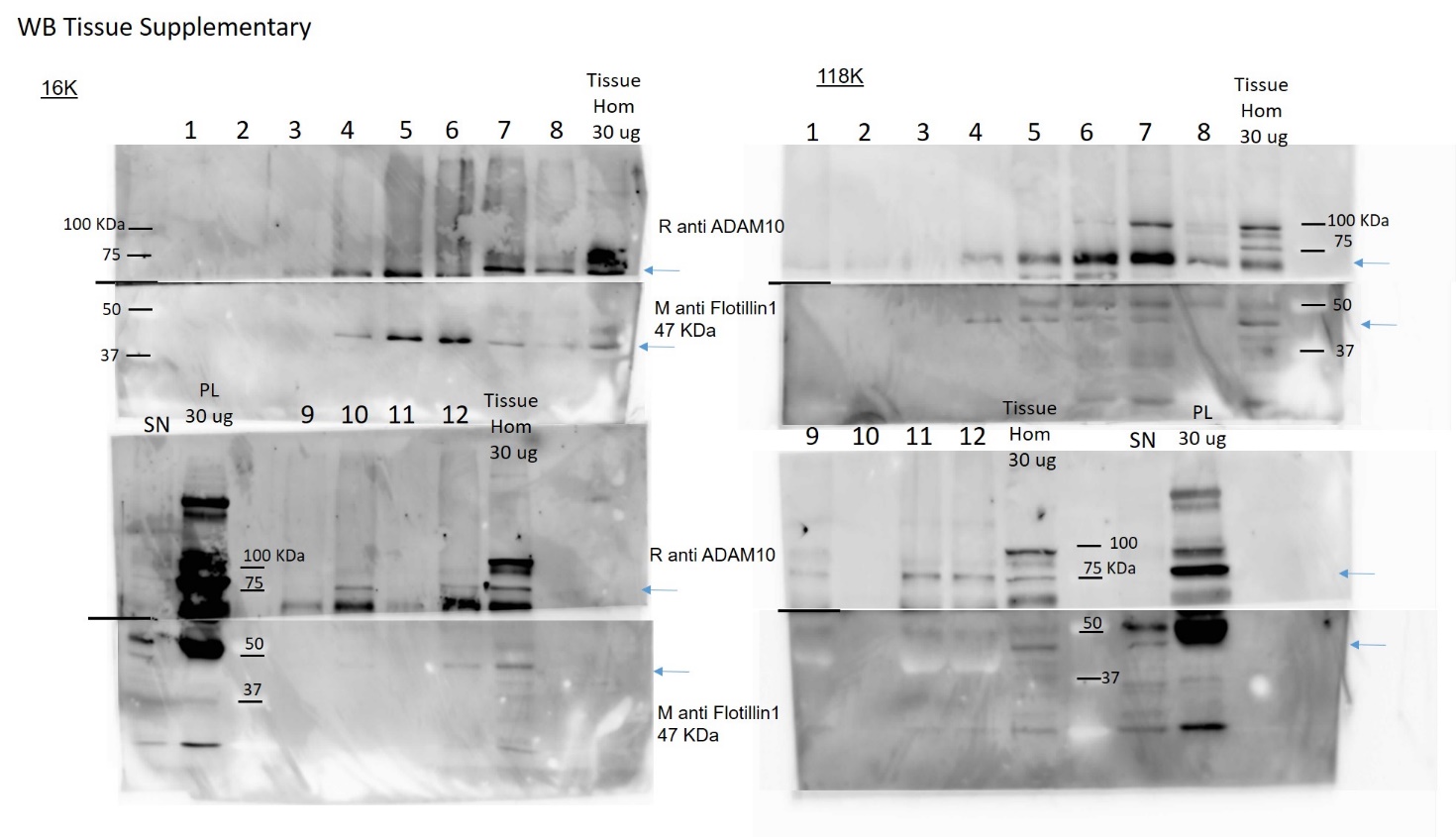

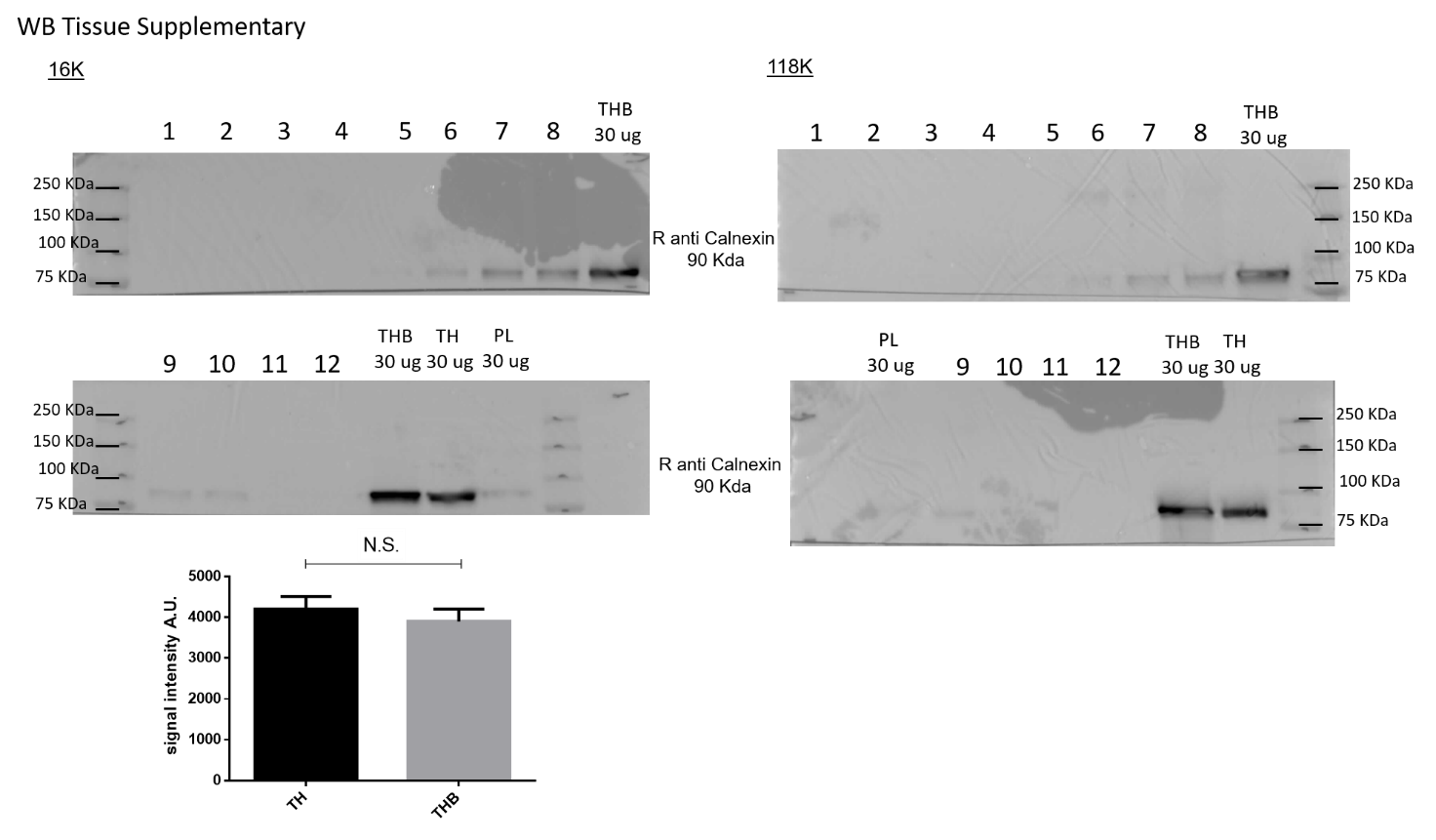
Fig.8SI. WB of Tissue 16k and 118k sample. DSG 1-12 sucrose fractions. PL: total plasma; TH: Skeletal muscle tissue homogenate after mechanical and enzymatic dissociation; THB: Skeletal muscle tissue homogenate before mechanical and enzymatic dissociation. Quantification of signal band intensity (Arbitrary unit- A.U.) of Calnexin in Tissue homogenate after mechanical and enzymatic treatment (TH) and Tissue homogenate before mechanical and enzymatic treatment (THB). Statistical analysis (Student t-test) indicate no significant differences between the samples analyzed.

**AFM analysis: Surface density calculation.**

Supplementary Table 1. Relative particle concentration of samples from multiple patients and enrichment procedures as estimated via AFM surface density (see Supplementary methods section). For all patients, samples enriched from tissue (TS) are more concentrated than those enriched from plasma (PL) following the same protocol (labeled here 16kG or 118kG). Relative concentrations are thus given here as TS:PL ratios. On average, TS samples contain around 100 times more particles than PL samples.

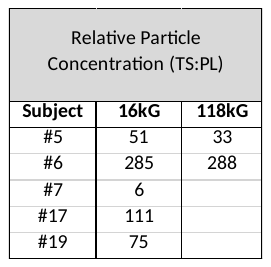

**DLS analyses:** autocorrelation function of the scattered intensity, with the fitting curve according to the Laplace Inversion via CONTIN algorithm; corresponding scattered intensity-weighted and number-weighted size distributions of scattering objects of each sample analyzed are available in Zenodo repository. https://doi.org/10.5281/zenodo.10579305

**CONAN assay**: results of the CONAN assay for each sample analyzed are available in Zenodo repository. https://doi.org/10.5281/zenodo.10579305

**NTA: Differences in Concentration and Size**

NTA was performed to assess particle concentration and size distribution of EVs isolated from TS and PL fractions obtained at 16kG and 118kG.

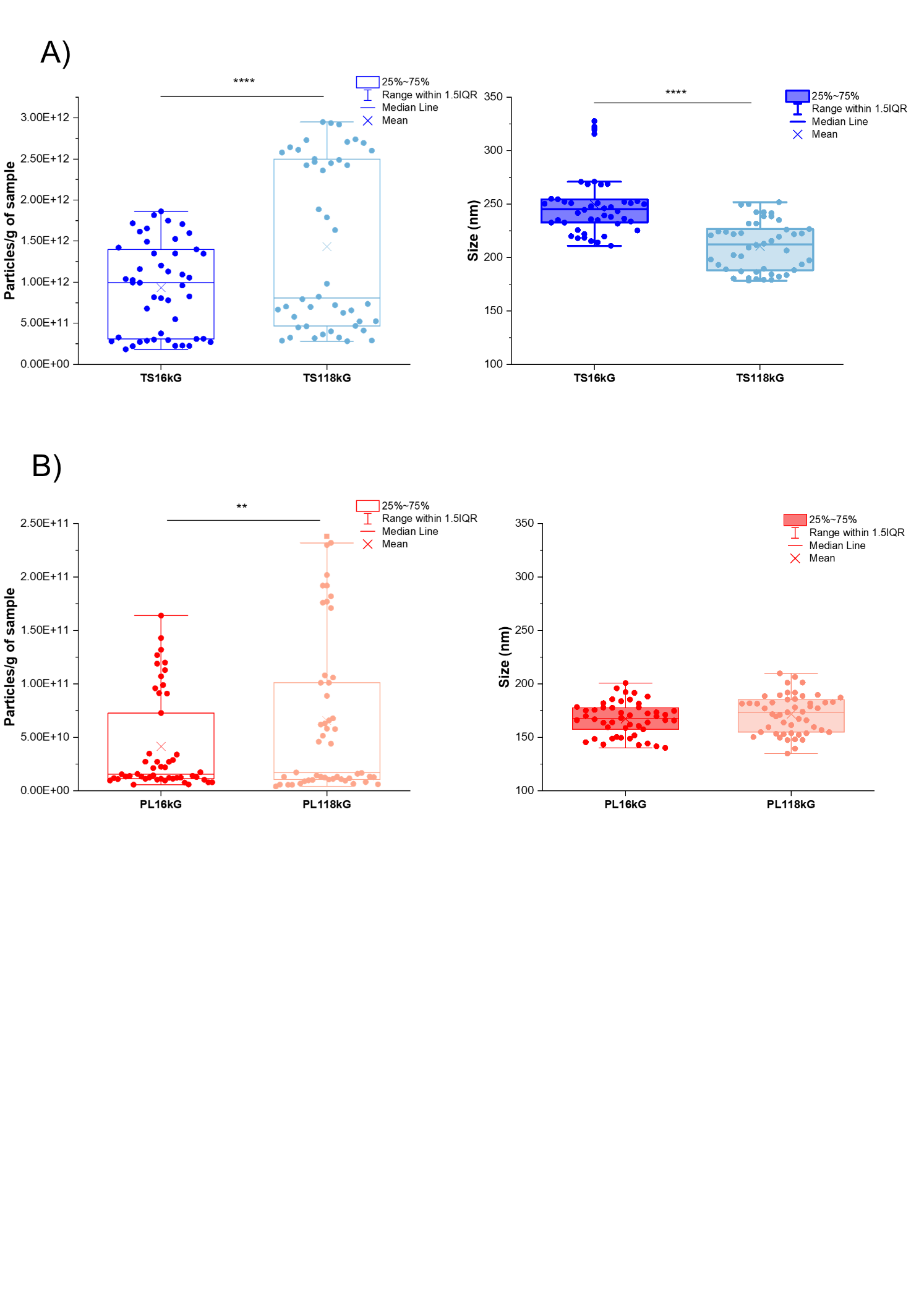

Figure9SI. Particle concentration and size distribution were measured by NTA. Particle concentration was normalized to starting material mass (1 g). Box plots represent the interquartile range (25th–75th percentile), with whiskers extending to 1.5×IQR; the central line indicates the median and the “X” denotes the mean. A) Particle concentration and mean size of TS EVs from 16kG and 118kG preparations; B) Particle concentration and mean size of PL EVs from 16kG and 118kG preparations.

Statistical significance was determined using Student’s t-test. P values: **p < 0.01, ***p < 0.001, ****p < 0.0001.

**References**

[1].	Lucien, F. *et al.* MIBlood-EV: Minimal information to enhance the quality and reproducibility of blood extracellular vesicle research. *Journal of Extracellular Vesicles.* **12**, 12385 (2023).

[2].	Grossi, I. *et al.* MicroRNA‑34a‑5p expression in the plasma and in its extracellular vesicle fractions in subjects with Parkinson’s disease: An exploratory study. *Int J Mol Med.* **47**, 533–546 (2021).

[3].	Bracht, J. W. P. *et al*. Platelet removal from human blood plasma improves detection of extracellular vesicle-associated miRNA. *J of Extracellular Vesicles*. **12**, 12302 (2023).

[4].	Coumans, F. A. W. *et al*. Methodological Guidelines to Study Extracellular Vesicles. Circ Res. **120**, 1632–1648 (2017).

[5].	Lacroix, R. *et al.* Standardization of pre-analytical variables in plasma microparticle determination: results of the International Society on Thrombosis and Haemostasis. *SSC Collaborative workshop*. *J Thromb Haemost.* (2013).

[6].	Radeghieri, A. *et al.* Active antithrombin glycoforms are selectively physiosorbed on plasma extracellular vesicles. *Journal of Extracellular. Biology* **1**, e57 (2022).

[7].	Paolini, L., Radeghieri, A., Civini, S., Caimi, L. & Ricotta, D. The epsilon hinge-ear region regulates membrane localization of the AP-4 complex. *Traffic.* **12**, 1604–1619 (2011).

[8].	Crescitelli, R., Lässer, C. & Lötvall, J. Isolation and characterization of extracellular vesicle subpopulations from tissues. *Nat Protoc.* **16**, 1548–1580 (2021).

[9].	Alvisi, G. *et al.* Intersectin goes nuclear: secret life of an endocytic protein. *Biochemical Journal.* **475**, 1455–1472 (2018).

[10]. Musicò, A. *et al.* Surface functionalization of extracellular vesicle nanoparticles with antibodies: a first study on the protein corona “variable”. *Nanoscale Advances.* **5**, 4703–4717 (2023).

[11]. Kowal, J. *et al.* Proteomic comparison defines novel markers to characterize heterogeneous populations of extracellular vesicle subtypes. *Proc Natl Acad Sci. U S A* **113**, E968-977 (2016).

[12]. Watanabe, S. *et al.* Skeletal muscle releases extracellular vesicles with distinct protein and microRNA signatures that function in the muscle microenvironment. *PNAS Nexus.* **1**, pgac173 (2022).

[13]. Théry, C. *et al.* Minimal information for studies of extracellular vesicles 2018 (MISEV2018): a position statement of the International Society for Extracellular Vesicles and update of the MISEV2014 guidelines. *J Extracell Vesicles.* **7**, 1535750 (2018).

[14]. Maiolo, D. *et al.* Colorimetric nanoplasmonic assay to determine purity and titrate extracellular vesicles. *Anal Chem.* **87**, 4168–4176 (2015).

[15]. Zendrini, A. *et al.* Augmented COlorimetric NANoplasmonic (CONAN) Method for Grading Purity and Determine Concentration of EV Microliter Volume Solutions. *Frontiers in Bioengineering and Biotechnology* **7**, (2020).

[16]. Ridolfi, A. *et al.* AFM-Based High-Throughput Nanomechanical Screening of Single Extracellular Vesicles. *Anal. Chem.* **92**, 10274–10282 (2020).

[17]. Ridolfi, A. *et al.* Particle profiling of EV-lipoprotein mixtures by AFM nanomechanical imaging. *Journal of Extracellular Vesicles.* **12**, 12349 (2023).

[18]. Nečas, D. & Klapetek, P. Gwyddion: an open-source software for SPM data analysis. *Open Physics.* **10**, 181–188 (2012).

[19]. Borup, A. *et al.* Comparison of separation methods for immunomodulatory extracellular vesicles from helminths. *Journal of Extracellular Biology.* **1**, e41 (2022).
